# REST/NRSF phosphorylation by CaMKIV regulates its transcriptional repressor activity and half-life

**DOI:** 10.1101/2025.09.04.674163

**Authors:** Hanako Tsushima Semini, Beatrice Corradi, Dmytro Shmal, Martina Chiacchiaretta, Nara Liessi, Andrea Di Fonzo, Emanuele Carminati, Andrea Armirotti, Anna Fassio, Luca Maragliano, Anna Rocchi, Fabio Benfenati

## Abstract

REST is a repressor of a large cluster of neural genes that homeostatically regulates neural activity. However, whether REST can be regulated by Ca^2+^ and Ca^2+^-activated kinases is unknown. We investigated Ca^2+^/calmodulin-dependent protein kinases (CaMKs) as upstream regulators of REST fate and activity. We show that REST is phosphorylated by CaMKIV at Ser-322 site located within the linker between the 5^th^-6^th^ Zn-finger domains. Phosphomimic REST mutant in Serine-322 decreased REST repressor activity and caused its transition from nucleus to cytosol, followed by degradation. Molecular dynamics simulations of the phosphomimic N-terminal REST and the DNA RE1 sequence revealed a sharp decrease in the stability of the REST-RE1 binding interface. Moreover, the homeostatic effects of CaMKIV on the amplitude of excitatory synaptic currents were inhibited by the genetic deletion of REST. The results demonstrate that CaMKIV phosphorylation has a crucial role in the homeostatic regulation of REST levels and repressor activity.

**Teaser:** CaMKIV phosphorylation homeostatically regulates the repressor activity of REST on neural genes and its half-life.

## INTRODUCTION

REST (RE1-Silencing Transcription Factor), also known as NRSF (Neuron-Restrictive Silencer Factor), is a transcriptional repressor that plays a key role in gene regulation (Bruce et al., 2004; Balls & Mandel, 2005). It contains 9 Zn-finger domains, through which it binds to a specific DNA sequence of 17-21 nucleotides named Repressor Element-1 (RE1; Mori et al., 1992; Kraner et al., 1992), at which it recruits co-factors (such as Co-REST, mSin3, and HDACs) to modify chromatin structure and repress the transcription of the downstream gene. RE1 elements have been identified in over 1,800 genes in the human and mouse genomes, displaying a variable number of copies and variable affinity for REST (Bruce et al., 2004). Many of the target genes of REST are neuronal genes, while others are involved in cell cycle regulation and cytoskeleton assembly. In non-neuronal tissues, REST is active and suppresses neuronal gene expression, while during neuronal differentiation, REST is downregulated, allowing neuronal genes to be expressed (Ballas et al., 2005; Yeo et al., 2009; Rodenas-Ruano et al., 2005; Aoki et al., 2011; see Baldelli & Meldolesi for review).

In addition to its role in neuronal differentiation, residual REST expression is preserved in mature neurons, albeit at low levels, and plays post-developmental roles in the brain. In neurons, REST is an essential actor of homeostatic plasticity responses. It is promptly induced by hyperactivity and migrates into the nucleus, where it inhibits the transcription of genes governing intrinsic excitability and synaptic transmission (Baldelli & Meldolesi, 2015; Michetti & Benfenati, 2024). Under conditions of neuronal hyperactivity, the induction of REST inhibits the expression of voltage-gated Na^+^ channels (Pozzi et al., 2013), blunts excitatory synaptic transmission by acting at the presynaptic level (Pecoraro-Bisogni et al., 2018), and boosts inhibitory synaptic transmission impinging onto excitatory neurons (Prestigio et al., 2021). REST is also involved in controlling changes in neuronal networks in response to early-life adversity, promoting resilience (Singh-Taylor et al., 2018; Mampay & Sheridan, 2019; Bolton et al., 2020). In addition, REST was also identified to have a key role in the regulation of lifespan and in stress resistance during aging by repressing hyperactivity (Lu et al., 2014; Zullo et al., 2019) and preventing senescence by regulating the autophagic flux (Rocchi et al., 2021). A homeostatic effect against hyperactivity was also observed in astrocytes, where REST increases glutamate uptake and potassium clearance from the synaptic environment (Centonze et al., 2023).

Beyond its canonical role in neurodevelopment and its pivotal role in regulating brain homeostasis, REST dysregulation has been implicated in a variety of pathological contexts, including epilepsy, neuroinflammation, brain ischemia, pain, Alzheimer’s and Huntington’s diseases, where it may contribute to neuronal vulnerability or adaptive responses (Buckley et al., 2010; Lu et al., 2014; McClelland et al., 2011, 2014; Hall et al., 2024; Carminati et al., 2021; Buffolo et al., 2021; Natali et al., 2023; Carminati et al., 2020; see Hwang & Zukin, 2018 for review).

REST shuttles between the nucleus, where it associates with a variety of interactors and actuates transcriptional repression, and the cytoplasm, where it remains inactive. Several regulators of REST favor its retention in the nucleus, such as the co-factors mSin3 and Co-REST and RILP (Shimojo & Hersh, 2006; Ballas & Mandel, 2005), and optogenetic disruption of these interactions inhibits REST activity in the nucleus (Paonessa et al., 2016). Although no full-length experimental structure is available for REST, domain mapping suggests that it is a modular protein comprising N-terminal and C-terminal repression domains and a central Zn-finger DNA-binding region. The latter adopts the canonical C2H2 architecture, consisting of nine conserved Cys2–His2 motifs arranged in tandem, each coordinating a Zn^2+^ ion and mediating DNA binding (Palm et al., 1998; Rockowitz et al., 2014; Brenner et al., 2005). Structural modeling and sequence alignment suggest that REST’s Zn-finger (ZNF) domains adopt the classical ββ-α fold, namely a β-hairpin followed by an α-helix, stabilized by Zn^2+^ coordination (Wolfe et al., 1999). Each Zn-finger module typically interacts with a tri-nucleotide segment of DNA, allowing REST to contact an extended nucleotide sequence and achieve high specificity (Klug, 2010).

REST activity is tightly regulated, not only at the transcriptional and epigenetic levels, but also through post-translational modifications, among which phosphorylation plays a central role. Indeed, phosphorylation was demonstrated to regulate REST half-life and turnover. Casein kinase 1 (CK1) was found to associate and phosphorylate REST at four serine residues (S^1013^, S^1024^, S^1027^, S^1030^) within two degron motifs in the C-terminal region, enabling recognition by the E3 ubiquitin ligase β-TrCP and targeting it for proteasomal degradation (Westbrook et al., 2008; Guardavaccaro et al., 2008; Kaneko et al., 2014). Moreover, extracellular signal-regulated kinases 1 and 2 (ERK1/2) were found to phosphorylate the proline-directed phosphorylation motifs in REST. ERK1/2 phosphorylation promotes rapid REST degradation, relieving repression of neuronal gene expression, while their dephosphorylation by the C-terminal domain small phosphatase 1 (CTDSP1) increases REST stability and half-life (Nesti et al., 2014). This ERK-mediated regulation plays an important role during neurogenesis, where ERK activation in response to growth factors leads to REST downregulation, allowing neuronal differentiation.

Despite these insights, the understanding of the full spectrum of REST phosphorylation sites and their functional consequences remains incompletely understood. While phosphorylation has been predominantly seen as a mechanism to downregulate REST in neural progenitors and differentiating neurons, it could also participate in regulating the downstream effects of REST overexpression during hyperactivity in mature neurons by negatively modulating REST transcriptional repression and returning REST levels to baseline. As calcium entry and calmodulin activation are the most likely signals triggering homeostatic plasticity and REST overexpression (Turrigiano, 2008, 2011), here we investigated whether REST can also be phosphorylated by Ca^2+^/calmodulin-dependent kinases (CaMKs) and whether its phosphorylation has any effect on its repressor activity, nuclear/cytosolic shuttling, and half-life.

We have found that, consistent with the presence of consensus phosphorylation sites for CaMKs, REST is phosphorylated by CaMKIV in primary neurons on a main site intercalated between the 5^th^-6^th^ Zn-finger domains, and its phosphorylation causes dissociation from the DNA RE1 motif, inhibition of its transcriptional repressor activity, escape from the nucleus to the cytoplasm where it undergoes degradation by extra-lysosomal proteases or pH-independent lysosomal proteases. Moreover, the homeostatic effects of CaMKIV on excitatory synaptic transmission were inhibited by the genetic deletion of REST. Overall, the effects of the CaMK-dependent phosphorylation of REST are important to blunt the homeostatic transcriptional responses of REST to hyperactivity and to return them to the basal condition, indicating a key physiological role of the interplay of CaMKIV/REST in homeostatic plasticity responses.

## RESULTS

### Inhibition of Ca^2+-^calmodulin-dependent kinases increases the levels of REST protein in primary neurons

Recent evidence suggests that post-translational modifications, including phosphorylation and ubiquitination, influence the stability and turnover of the REST protein. ERK1/2 and CK1 kinases phosphorylate REST and promote its degradation by the ubiquitin proteasome system (Westbrook et al., 2008; Kaneko et al., 2014; Nesti et al., 2014). Given that REST is upregulated in mature neurons by hyperactivity, a condition in which Ca^2+^ entry occurs, we tested the possibility that phosphorylation by CaMKs regulates REST turnover and fate in neurons. We first evaluated whether Ca^2+^/calmodulin (CaM) signaling interferes with REST stability. When primary cortical neurons were incubated with the CaM inhibitor W7, an increase in REST protein was observed (**Figure 1A**).

**Figure 1.**
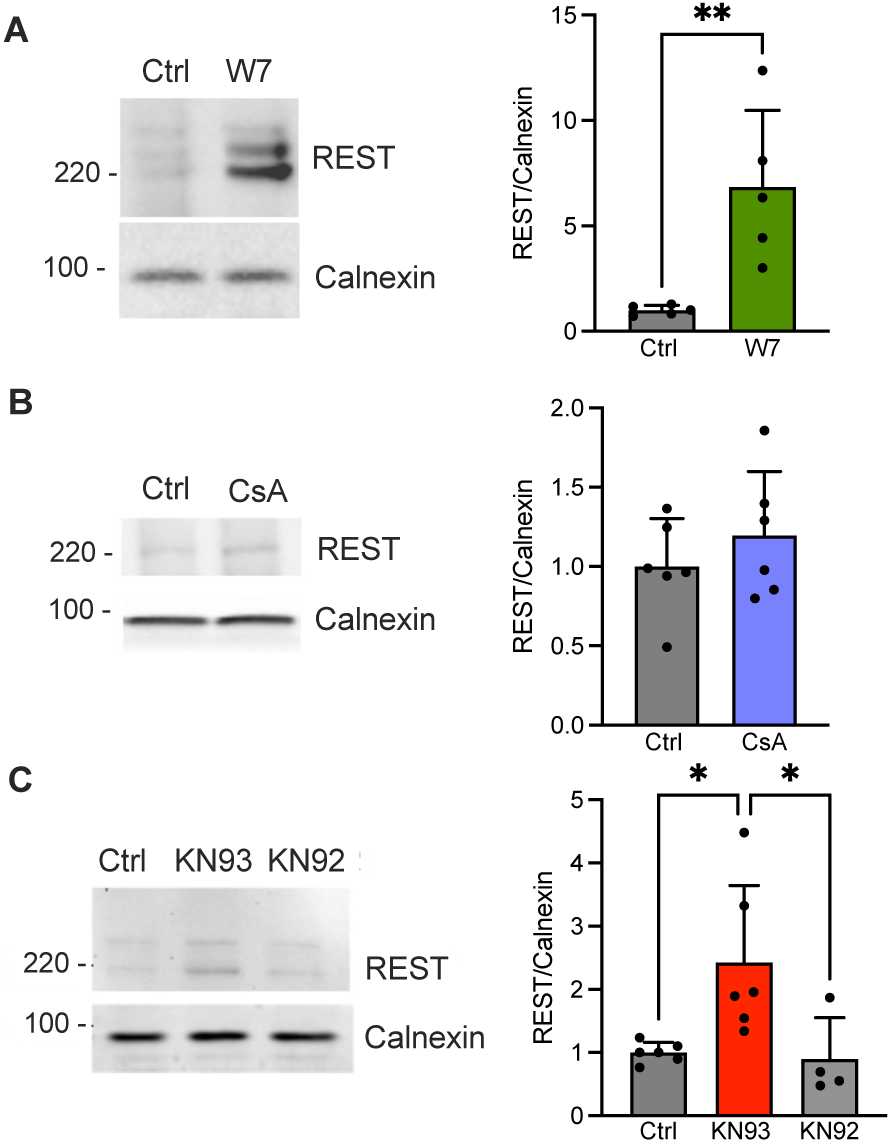
Inhibition of Ca^2+^-calmodulin-dependent activities increases the endogenous levels of REST protein in primary mouse cortical neurons. **A.** *Left:* Representative REST immunoblots of primary cortical neurons (DIV14) treated with the cell-permeable competitive Ca^2+^-calmodulin antagonist W7 (20 µM). Calnexin was used as a control for equal loading. *Right:* Mean (± sem with individual experimental points) REST protein expression shown as a ratio between REST and calnexin immunoreactivities. **p=0.0071; unpaired Student’s *t*-test (n=5 independent neuronal preparations). **B.** *Left:* Representative REST immunoblots of DIV 14 primary cortical neurons treated with the phosphatase inhibitor cyclosporin A (CsA; 1 µM). Calnexin was used as a control for equal loading. *Right:* Mean (± sem with individual experimental points) REST expression shown as a ratio between REST and calnexin immunoreactivities. p=0.36; unpaired Student’s *t*-test (n=5 independent neuronal preparations). **C.** *Left:* Representative REST immunoblots of DIV14 primary cortical neurons treated with vehicle (Ctrl), the cell-permeable CaMK inhibitor KN93 (10 µM), or its inactive derivative KN92 (10 µM). Calnexin was used as a control for equal loading. *Right:* Mean (± sem with individual experimental points) REST protein expression shown as a ratio between REST and calnexin immunoreactivities for the three treatments. *p=0.031, Ctrl *vs* KN93; *p=0.032, KN93 *vs* KN92; one-way ANOVA/Holm-Sidak’s tests (n=5 independent neuronal preparations).

Since CaM activates various kinases and phosphatases, we asked which downstream effector could be involved. Primary cortical neurons were treated with either the calcineurin inhibitor ciclosporin A (CsA) or with the competitive CaMK blocker KN93. While inhibition of calcineurin did not affect REST protein levels (**Figure 1B**), KN93, at concentrations inhibiting all CaMKs, induced a significant increase in REST protein levels that was completely absent when its inactive analogue KN92 was used (**Figure 1C**). None of the treatments was associated with any change in REST mRNA levels, as shown by RT-qPCR analysis (**Figure S1**). The same treatments, performed on primary cortical astrocytes, did not alter the levels of both REST protein and mRNA (**Figure S2**), indicating that the effects of CaMK inhibition were specific for neurons.

### Expression of constitutively active CaMKI and CaMKIV decreases the expression and the transcriptional repressor activity of REST in HEK293T cells

KN93 is a broad-spectrum CaMK inhibitor with several molecular targets, and thus, the results do not identify the specific CaMKs that mediate the effects on REST protein levels. To address this point, we overexpressed CaMKI, CaMKII, and CaMKIV as the wild-type (WT) form, and as constitutively active (CA) and kinase-dead (KD) variants in HEK293T cells that constitutively express REST. Under these experimental conditions, we evaluated (i) the changes in the expression levels of the REST protein by immunoblotting as a ratio between REST and calnexin immunoreactivities; and the effects on the REST repressor activity by using a gene reporter assay in which *Renilla* luciferase is driven by the SV40 promoter fused to a triple RE1*cis* site from the promoter region of the *SCN2A* gene, a well-known REST target (**Figure 2**; Paonessa et al., 2016; see Materials and Methods).

**Figure 2.**
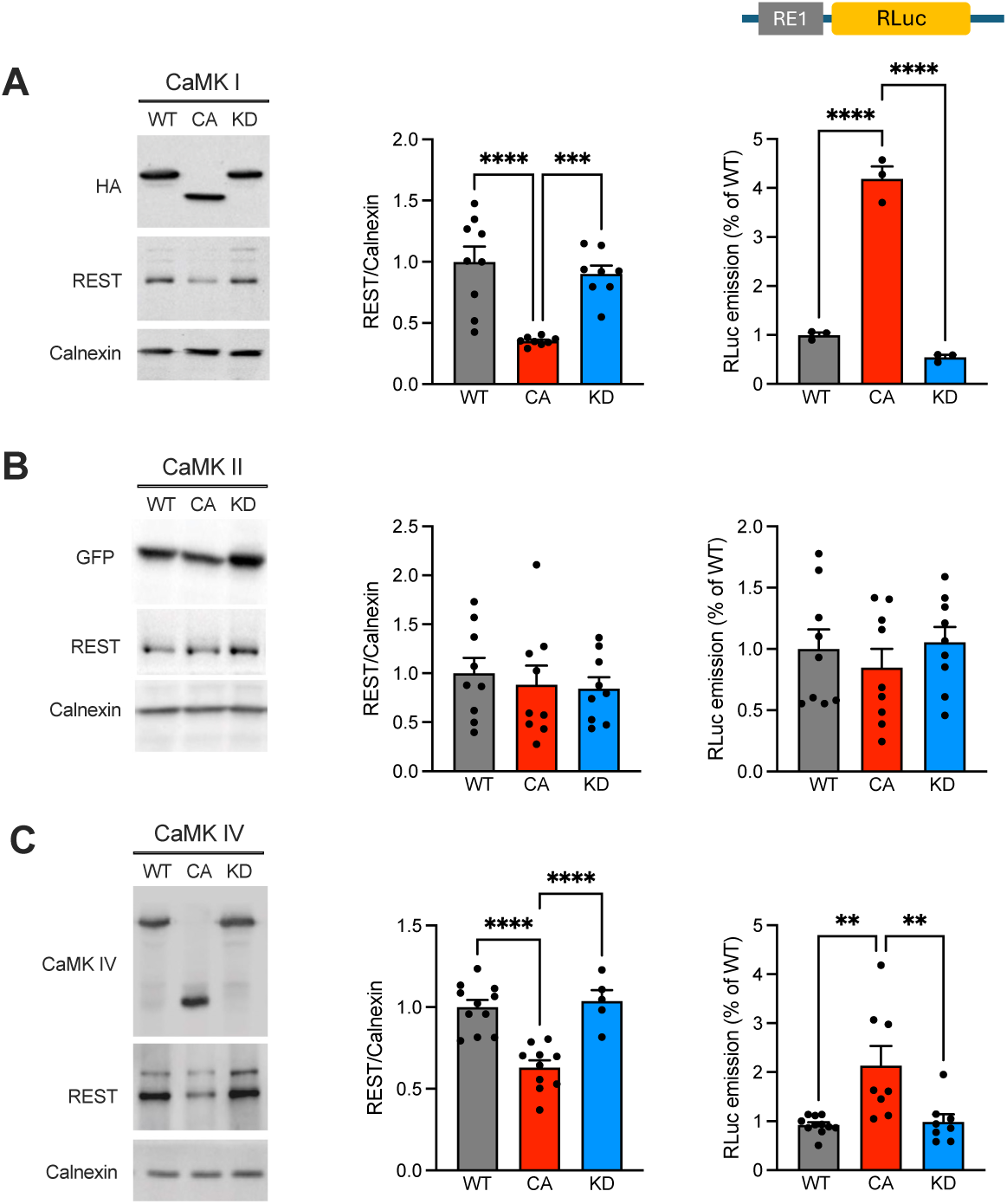
Expression of wild-type, constitutively active, or kinase-dead CaM kinases in HEK293T cells implicates CaMKI and CaMKIV in the control of protein levels and transcriptional repressor activity of REST. **A-C.** *Left:* Representative immunoblots monitoring the expression of wild-type (WT), constitutively active (CA), and kinase-dead (KD) CaMKs, REST, and calnexin, respectively, in HEK293T cells, 48 h after transfection with the kinase constructs. The expression of CaMKI (**A**) and CaMKII (**B**) was followed using antibodies to HA and GFP fused tags, respectively, while CaMKIV (**C**) was detected with anti-CaMKIV-specific antibodies. *Middle:* Mean (± sem with individual experimental points) REST protein levels after expression of the three variants of CaMKI (**A**), CaMKII (**B**), and CaMKIV (**C**), shown as REST/calnexin immunoreactivity ratios. *Right:* A RE1-Luciferase construct, with *Renilla* Luciferase (*RLuc*) gene cloned downstream of the RE1 sequence, was used to detect the repressor activity of endogenous REST in HEK293T cells, transfected with the three variants of CaMKI (**A**), CaMKII (**B**), and CaMKIV (**C**). The *Renilla/Firefly* Luciferase chemiluminescence ratio is shown as mean (± sem with individual experimental points). Panel A - REST/calnexin: ***p=0.0007 CA *vs* KD, ****p<0.0001 CA *vs* WT; *RLuc*: ****p<0.0001 CA *vs* WT or KD. Panel B - REST/calnexin: p=0.86 CA *vs* WT, p=0.98 CA *vs* KD; *RLuc*: p=0.74 CA *vs* WT, p=0.58 CA *vs* KD. Panel C - REST/calnexin: ****p<0.0001 CA *vs* WT or KD; *RLuc*: **p=0.0018 CA *vs* WT; **p=0.0057 CA *vs* KD; one-way ANOVA/Tukey’s tests (n=8-9 independent HEK293T cell preparations).

Western blotting analysis revealed that the constitutively active forms of both CaMKI and CaMKIV significantly decreased REST levels, while the constitutively active form of CaMKII was virtually ineffective (**Figure 2A-C**, *left and middle panels*). No effects with respect to the WT forms of the kinases were observed with the respective kinase-dead variants, suggesting that the endogenous activation of these kinases is not functional in this cell line under these conditions. In agreement with the observed decrease in REST levels, we observed a markedly increased luciferase signal in HEK293T cells transfected with the constitutively active forms of both CaMKI and CaMKIV and no effect with the active form of CaMKII (**Figure 2A-C**, *right panels*), indicating a corresponding drop in the REST repressor activity at RE1 sites. The data indicate that CaMKI and CaMKIV are potentially implicated in the regulation of REST protein stability, at least in HEK293Tcells.

### Knockdown of CaMKIV, but not of CaMKI, in primary neurons recapitulates the effects of CaMK inhibition on REST protein levels

To get insights into the mechanism underlying the CaMKI/CaMKIV-mediated regulation of REST levels in neurons, the REST protein level and activity were analyzed after silencing of either CaMKI or CaMKIV by RNA interference using lentiviral vectors (**Figure 3** and **Figure S3**). While we used a very effective, previously reported shRNA for CaMKII (Ageta-Ishihara et al., 2009), we tested two distinct shRNAs (sh#1 and sh#2) for their efficiency in knocking down CaMKIV (**Figure S3A**). Both shRNAs effectively downregulated CaMKIV in primary neurons with >60% decrease in CaMKIV levels. We selected sh#2 for the successive experiments and first tested the specificity of the knockdown with respect to the other CaMKs and REST. The shRNA for CaMKIV was very selective and was not associated with any change in REST, CaMKI, or CaMKII mRNA levels (**Figure S3B-D**).

**Figure 3.**
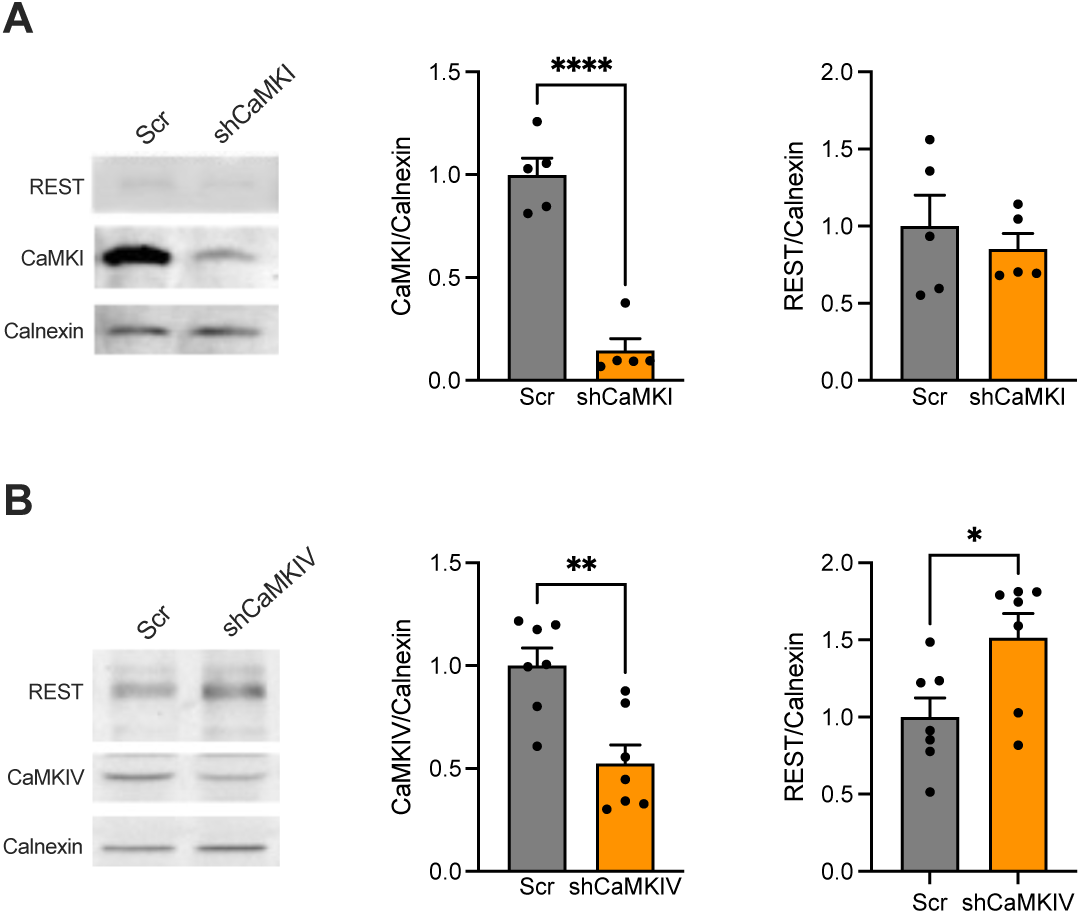
Downregulation of CaMKIV, but not of CaMKI, increases the endogenous level of REST in mouse primary cortical neurons. **A.** Primary mouse cortical neurons were transduced at DIV 7 with lentiviruses encoding shRNAs to downregulate CaMKIα (shCaMKI) or a scrambled version thereof (shSCR). *Left:* Representative immunoblots monitoring the expression of endogenous CaMKIα, REST, and calnexin in the neuronal preparations at DIV14. *Middle and Right:* Mean (± sem with individual experimental points) CaMKIα (*middle*) and REST (*right*) expression shown as ratios with calnexin immunoreactivity. CaMKIα: ****p<0.0001; REST/NRSF: p=0.53; unpaired Student’s *t*-test (n=5 independent preparations). **B.** Primary mouse cortical neurons were transduced at DIV 7 with lentiviruses encoding shRNAs to knock down CaMKIV (shCaMKIV) or a scrambled version thereof (shSCR). *Left:* Representative immunoblots monitoring the expression of endogenous CaMKIV, REST, and calnexin in the neuronal preparations at DIV14. *Middle and Right:* Mean (± sem with individual experimental points) CaMKIV (*middle*) and REST (*right*) expression, shown as ratio with calnexin immunoreactivity. CaMKIV: **p=0.0024; REST/NRSF: *p=0.025; unpaired Student’s *t*-test (n=7 independent preparations).

To ascertain whether both CaMKI and CaMKIV are active in decreasing REST levels in neurons, we silenced either CaMKI or CaMKIV by transducing neurons with lentivirus encoding the respective shRNAs. Western blotting analysis revealed that silencing CaMKIV, but not CaMKI, induced an increase in REST protein levels (**Figure 3A,B**). In conclusion, the data suggest that CaMKIV is responsible for the post-transcriptional modulation of REST protein levels in primary neurons, which is in line with the preliminary results obtained with the CaMK inhibitors. Notably, while CaMKI is expressed by neurons and astrocytes, CaMKII and CaMKIV are virtually neuron-specific (**Figure S4**), indirectly confirming the role played by CaMKIV in the regulation of REST levels that was observed in neurons but not astrocytes under the same experimental conditions. Moreover, the distinct effects of CaMKI and CaMKIV in neurons may depend on their subcellular distribution. Whereas CaMKIV is predominantly nuclear, and therefore positioned to directly interact with nuclear REST, CaMKI is largely cytosolic, which limits its ability to efficiently target REST in neurons despite being effective in heterologous systems such as HEK293T.

### REST degradation is triggered by CaMKIV phosphorylation

Phosphorylation by CK1 and ERK1/2 has been shown to target REST to the ubiquitin-proteasome degradation. Thus, we sought to investigate whether phosphorylation by CaMKIV, also decreasing the levels of REST protein, was associated with a similar mechanism. We initially considered three possible pathways of degradation, namely the proteosomal, macroautophagic, lysosomal, or cytosolic proteolytic degradation (**Figure 4A**).

**Figure 4.**
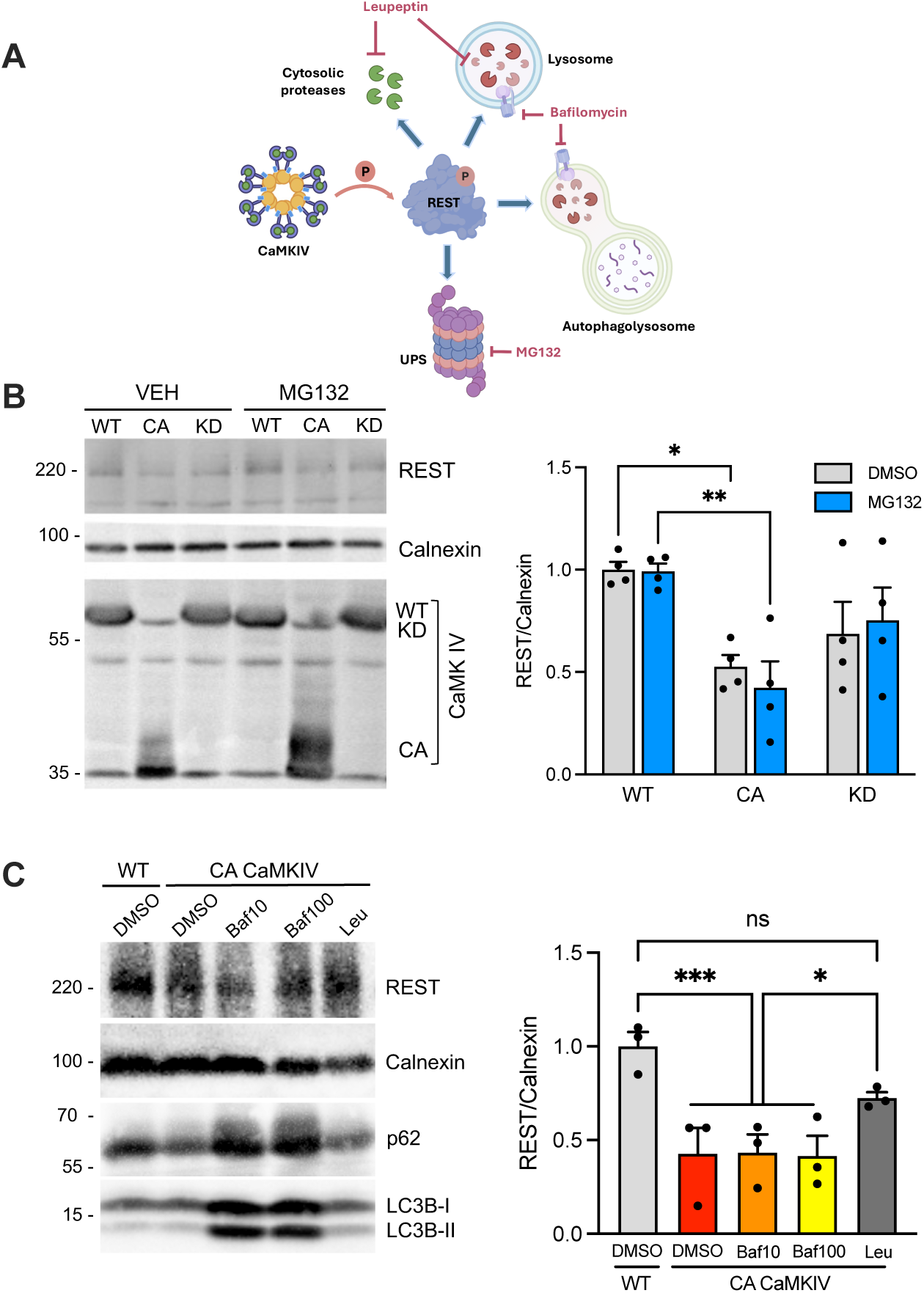
CaMKIV phosphorylation of REST triggers its lysosomal degradation. **A.** Schematics of the four possible degradation pathways that CaMKIV-phosphorylated REST can undertake and the three inhibitors that were used to dissect the fate of REST. UPS, ubiquitin proteasome system. **B.** HEK293T cells were transfected with WT, constitutively active (CA), or kinase dead (KD) CaMKIV and treated with cycloheximide (CHX) to inhibit further translation. Then, cells were treated for 16 h with either vehicle (DMSO) or MG132 (5 µM), an inhibitor of the ubiquitin-proteasome system. *Left:* Representative immunoblots monitoring the expression of wild-type (WT), constitutively active (CA), and kinase-dead (KD) CaMKs, REST, and calnexin, respectively, in HEK293T cells, 48 h after transfection with the kinase constructs. *Right:* Mean (± sem with individual experimental points) REST protein levels, shown as ratios with calnexin immunoreactivity under the various experimental conditions. No effects of inhibition of the ubiquitin-proteasome system were observed. *p=0.017, CA *vs* WT in DMSO; **p=0.0048, CA *vs* WT in MG132); 2-way ANOVA/Tukey’s tests (n=4 independent preparations). **C.** HEK293T cells were transfected with either WT or constitutively active CaMKIV (CA CaMKIV) and treated with cycloheximide to inhibit further translation. Then, cells were treated for 16 h with vehicle (DMSO), bafilomycin A1 (Baf, 10 and 100 nM; an inhibitor of lysosomal acidification and autophagy), and leupeptin (Leu, 40 µM; an inhibitor of the lysosomal and cytosolic proteases). *Left:* Representative immunoblots monitoring the expression of REST, calnexin, and the autophagy markers p62 and LC3B-II. *Right:* Mean (± sem with individual experimental points) of REST protein levels, shown as ratios with calnexin immunoreactivity under the various experimental conditions. ***p=0.0010, CA *vs* WT in DMSO and WT/DMSO *vs* CA/Baf10 and 100; *p=0.0264 CA/Leu *vs* CA/DMSO; 2-way ANOVA/Tukey’s tests (n=4 independent preparations).

To this aim, we transfected HEK293T cells with WT, constitutively active, or kinase-dead CaMKIV and treated them with cycloheximide (CHX) to block REST neo-synthesis. Then, we treated the cells with either vehicle or MG-132, a specific inhibitor of the ubiquitin-proteasome system. As reported above (see Figure 2), only cells expressing CA-CaMKIV exhibited significantly decreased REST levels. However, this downregulation of REST protein was not attributable to proteasome-mediated degradation, as MG-132 was totally ineffective in antagonizing the decreased REST levels associated with the expression of the constitutively active kinase (**Figure 4B**). The absence of effects of MG-132 in cells expressing WT- or KD-CaMKIV also indicates that, in the absence of specific activation of signal transduction pathways, REST is not constitutively targeted to the proteasome.

Next, we addressed the possibility that REST is degraded by alternative pathways, such as in autophagolysosomes or lysosomes. To ascertain this, we transfected HEK293T cells with either WT- or CA-CaMKIV and blocked REST neo-synthesis (see above) before treating the cells with vehicle, bafilomycin A1 (an inhibitor of v-ATPase responsible for lysosomal acidification), or leupeptin (an inhibitor of lysosomal and cytosolic proteases). The differential effect of these two blockers is demonstrated by the significant increase in the autophagy-related proteins LC3B-II and p62 observed after treatments with bafilomycin A1, but not with leupeptin (**Figure 4C**, *left panel*; **Figure S5A,B**). The results show that only leupeptin could reverse the drop in REST protein levels associated with expression of CA-CaMKIV (**Figure 4C**), indicating that REST degradation is likely mediated by proteases that are either cytosolic or not impaired by the loss of lysosomal acidification.

### REST is phosphorylated by CaMKIV on Serine322 between the 5^th^ and the 6^th^ Zn-finger domain

CaMKIV, together with CaMKI and CaMKII, is collectively defined as a “multifunctional CaM kinase” for its ability to phosphorylate a wide range of substrates (Xu et al., 2024; Kaiser et al., 2024). CaMKIV needs to be activated in the cytoplasm by CaMK kinase (CaMKK) before entering the nucleus, where it acts as a potent activator of Ca^2+^-dependent transcriptional effects through phosphorylation of downstream transcriptional effectors, such as CREB, MeCP2, and HDAC4 (Lemrow et al., 2004; Matthews et al., 1994; Xu et al., 2024). We thus searched for the presence of putative consensus sequences for CaMKIV phosphorylation in the primary structure of mouse and human REST (**Figure 5A**; White et al., 1998).

**Figure 5.**
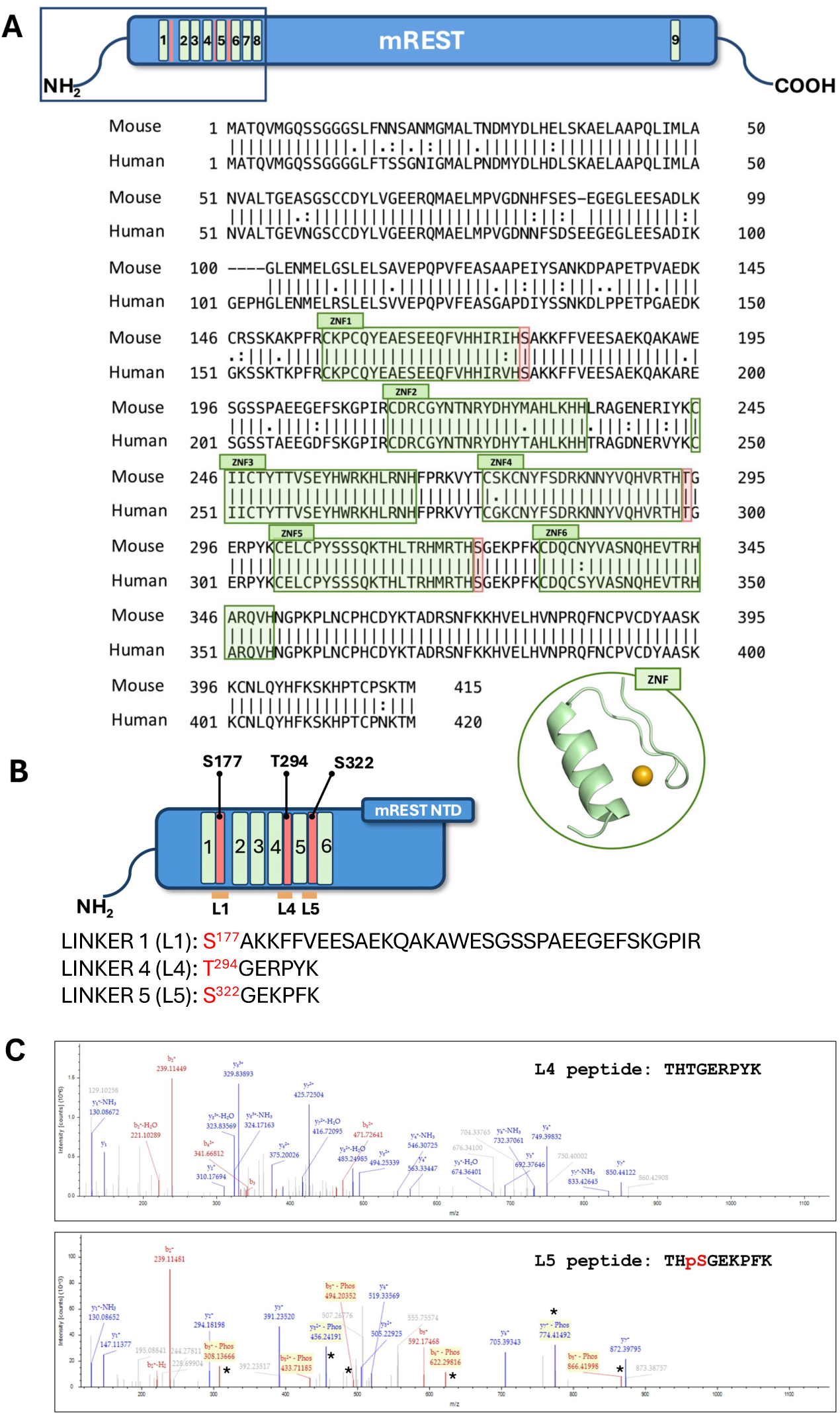
Serine-322 in the linker between the 5^th^-6^th^ Zn-finger domains is the major CaMKIV phosphorylation site in mouse REST. **A.** Alignment of the primary structures of human, rat, and mouse REST. Amino acids are shown in single-letter code. Sequences shaded in green refer to the Zn-finger domains (numbered 1 to 6), while three putative phosphorylation sites (S^177/182^, T^294/299^, and S^322/327^ in mouse and human, respectively) that match the *consensus* sequence for CaMKs (S/T-XX-R/K) and are conserved between mouse and human REST are highlighted in red. **B.** The S^177^, T^294^, and S^322^ phosphorylation sites encompass the linker peptides between the 1^st^-2^nd^ (L1), the 4^th^-5^th^ (L4), and the 5^th^-6^th^ (L5) in mouse REST Zn-finger domains, respectively. **C.** Tandem mass spectra for L4 (top) and L5 (bottom) tryptic peptides derived from HA-tagged recombinant mouse REST expressed in HEK293T cells, affinity-purified with anti-HA magnetic beads, and phosphorylated on beads by purified constitutively active CaMKIV. L5 shows phosphorylation of S322, as clearly demonstrated by the corresponding diagnostic sequence fragment ions (asterisk in the spectrum).

We identified three putative CaMKIV phosphorylation sites within the 8 ZNF central cluster region of mouse and human REST, namely Serine177 (S177), Threonine294 (T294), and Serine322 (S322), located at 1-position of the ZNF linkers L1, L4, and L5. All three sites display the required −3 Arginine residue but have the required hydrophobic residue at −4, rather than at the canonical −5 position (where they have a basic Histidine). While all three putative sites have a hydrophobic residue at +4 that is highly preferred by CaMKI, T294, and S322 are particularly interesting since they display a +2 acidic residue (Glutamate) found in optimal consensus sequences for CaMKII. In contrast, S177 has a basic residue instead. The three putative phosphorylation sites were analyzed for their phosphorylation potential for CaMKs using the prediction software Musite and NetPhos3.1. As shown in **Table S1**, in both human (hREST) and mouse (mREST) sequences, the S322 site has the highest phosphorylation potential, while S177 displays the lowest. In addition to these structural issues, the REST peptide linkers encompassing the T294 and S322 putative sites share a strong analogy with other transcription factors containing ZNF clusters, such as the transcription factor Ikaros (Jantz & Berg, 2004), in which ZNF domains are usually organized in arrays joined by short 5-7-mer linker sequences that often start with a phosphorylatable residue. In the REST protein, the linker peptide containing the putative site 1 (S177) is the least conserved from mouse to man and does not have any resemblance to the short linker peptide sequences joining adjacent ZNF domains, being a long 36-mer peptide connecting ZNF1 with ZNF2. On the contrary, the 7-mer T294- and S322-containing linkers L4 and L5 have a strong resemblance to Icaros ZNF linkers. For all these reasons, we concentrated our attention on T294 and S322 and tested whether they can be phosphorylated by CaMKIV (**Figure 5B**).

To this aim, we expressed Myc-tagged recombinant mREST in HEK293T cells, affinity-purified it with anti-Myc magnetic beads, and phosphorylated it on beads by purified constitutively active CaMKIV. Purified phosphorylated mREST was subjected to tryptic cleavage and tandem mass spectrometry (MS-MS). Tandem mass spectra for L4 (top) and L5 (bottom) tryptic peptides revealed phosphorylation of S322, but not of T294, as clearly demonstrated by the corresponding diagnostic sequence fragment ions (**Figure 5C**).

### Phosphomimic mutants of REST translocate to the cytoplasm, where they undergo degradation

The location of T294 and S322 in the very short L4/L5 ZNF linkers of mREST resembles that of the transcription factor Ikaros, in which phosphorylation of the first residue in the short ZNF linkers impairs the binding of the transcription factor to DNA (Jantz & Berg, 2004). This raises the possibility that phosphorylation of S322 (and, possibly, of T294) affects the interaction between REST and the RE1 motifs in the DNA. To verify that, we replaced either or both phosphorylatable residues with negatively charged Glutamate to generate phosphomimic REST mutants and examine their activity on REST nucleus/cytoplasm shuttling, fate, and transcriptional repression (**Figure 6**). HEK293T cells were transfected with either WT mREST or with the single (T294E, TE, and S322E, SE) or the double (TESE) mREST pseudo-phosphorylated mutants, using cytosolic mCherry and transfection reporter and Hoechst staining to morphologically identify the nuclear areas (**Figure 6A**, *left panel*). Morphometric analysis of the presence of REST immunoreactivity within the nuclear area was performed by calculating the Manders’ overlap coefficient, which showed a significant degree of the extent of colocalization for the SE mutant and smaller decreases for the other mutants, suggesting that the SE phosphomimic mutant of mREST has a lower affinity for its DNA target sequences and partially escapes from the nucleus (**Figure 6A**, *right panel*).

**Figure 6.**
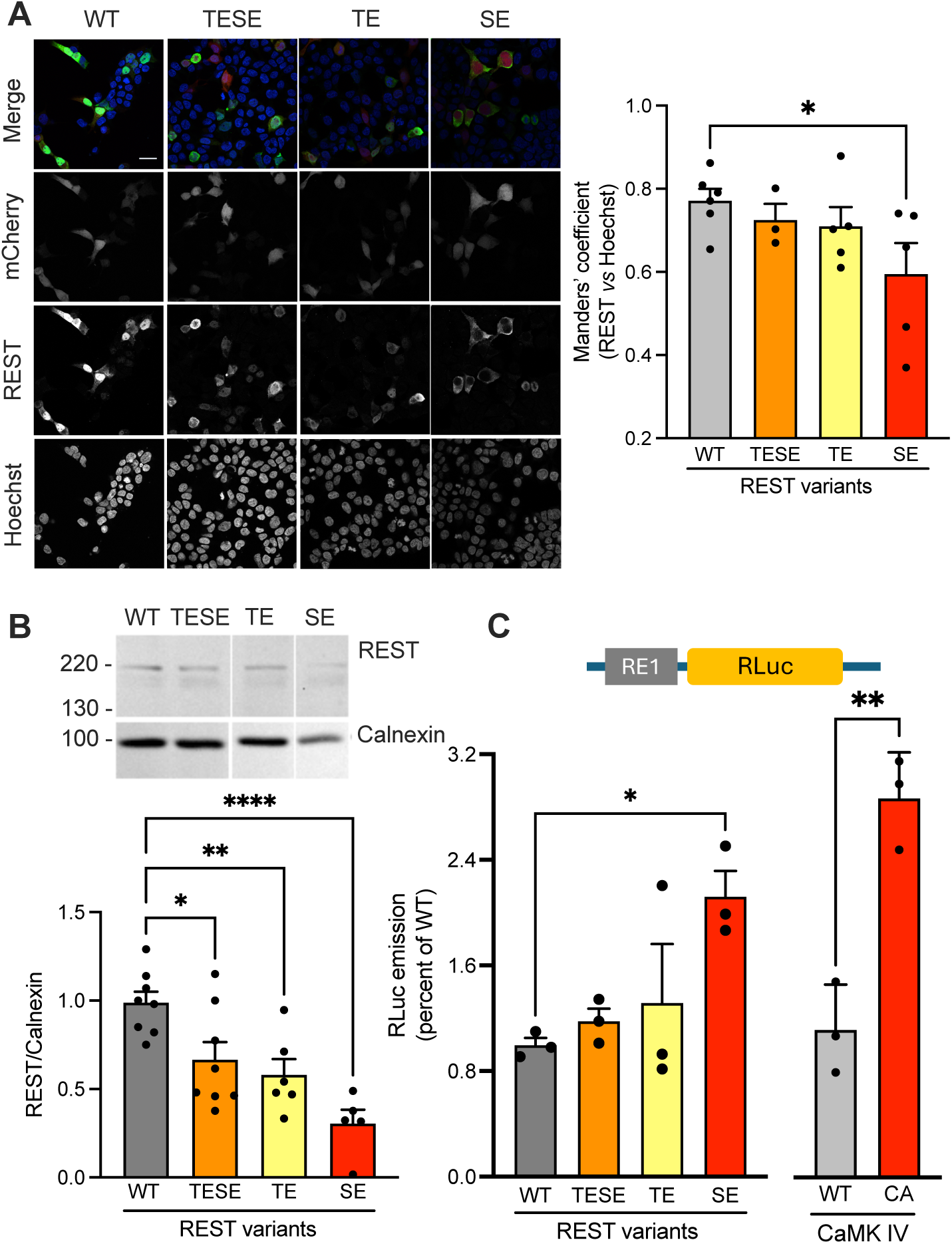
The phosphomimic S322E REST mutant has an impaired repressor activity and shuttles to the cytosol, where it undergoes degradation in HEK293T cells. **A.** HEK293T cells were transfected with WT REST or with its T294E (TE), S322E (SE), and S322E/T294E (TESE) phosphomimic mutants with soluble mCherry as transfection reporter. *Left:* Representative samples for the 4 experimental groups are shown, in which REST immunostaining, Hoechst nuclear staining, and mCherry fluorescence were used to evaluate the cytosolic/nuclear localization of REST mutants. Right: Mean (± sem with individual experimental points) of the Manders’ coefficient evaluating the extent of overlap between REST immunoreactivity and Hoechst nuclear staining. *p=0.039; one-way ANOVA/Dunnett’s tests. **B.** A RE1-Luciferase construct, with *Renilla* luciferase (*RLuc*) gene cloned downstream the promoter region containing triple RE1 sequence from *Scn2a*, was used to assess the extent of transcriptional repression exerted by the expression of WT REST or of its phosphomimic mutants in HEK293T cells. *Left:* The *Renilla/Firefly* Luciferase chemiluminescence ratio is shown as mean (± sem with individual experimental points). *Right:* As a positive control, either WT or catalytically active (CA) CaMKIV was transfected in the same HEK293T cells transfected with WT mREST. *p=0.032; one-way ANOVA/Dunnett’s tests. *p=0.0035; unpaired Student’s *t*-test (n=3 independent experiments). **C.** HEK293T cells were transfected with WT REST or with its T294E (TE), S322E (SE), and S322E/T294E (TESE) phosphomimic mutants. The levels of phosphomimic SE are strongly decreased in comparison to those of WT REST, while phosphomimic TE and the double phosphomimic TESE exhibit smaller decreases. *p=0.022, **p=0.0072, ****p<0.0001; one-way ANOVA/Dunnett’s tests. (n=5-8 independent experiments).

The successive analysis aimed at evaluating the total REST protein levels after transfection of HEK293T cells with either WT mREST or with the single or double mREST pseudo-phosphorylated mutants revealed that the decreased nuclear localization of REST was paralleled by an increased cytoplasmic degradation that was particularly dramatic for the SE mutant (**Figure 6B**).

Finally, we analyzed REST repressor activity after transfection of HEK293T cells with the three phosphomimic mutants and compared it with WT mREST in the absence or presence of co-transfected CA-CaMKIV by using the gene reporter assay in which *RLuc* is driven by the SV40 promoter fused to a triple RE1*cis* site. Strikingly, *RLuc* expression was significantly increased with the SE REST mutant, mimicking the decrease in transcriptional repression observed with CA-CaMKIV (**Figure 6C**). Taken together, the data on the REST phosphomimic mutant SE indicate that phospho-S322 is the phosphorylation site that mimics the effects of CA-CaMKIV overexpression on REST localization, stability, and repressor activity.

### Phosphomimic substitutions destabilize the mREST:RE1 interaction interface and promote local structural collapse

To evaluate the impact of phosphomimic substitutions on the stability of the mREST:RE1 interaction interface, we conducted atomistic molecular dynamics (MD) simulations on both wild-type and mutant complexes. We focused on the mREST segment V130-H267. The mREST:RE1 complex was reconstructed using a multi-step protocol that included structural modeling, protein-DNA docking, complex refinement, and optimization of Zn^2+^ coordination (**Figures S6-S11**). The resulting structure was used as the starting conformation for three independent 1-µs replicate MD simulations (indicated with R1 to R3).

To monitor global, structural deviations over time, we computed the Root Mean Square Deviation (RMSD) using all the heavy atoms of the protein and of the DNA. The RMSD profiles are stable around values between 2.8 and 4 Å, indicating no significant structural deviations from the starting configuration (**Figure S12**). We quantified the integrity of the protein-DNA interface by measuring the distance between a selected phosphate atom of the DNA backbone, adjacent to the first Zn-finger module, and the center of mass (COM) of the Zn-coordinating side chain atoms (Cysteine and Histidine residues) as a distance-based descriptor. This distance was stable throughout all simulations of the WT system, with average values of 10.3 ± 2.4 Å (R1), 11.1 ± 1.5 Å (R2), and 7.6 ± 0.7 Å (R3), and no local structural deviations were observed (**Figure 7A**), indicating that the WT mREST:RE1 complex maintains a stable interface over time.

**Figure 7.**
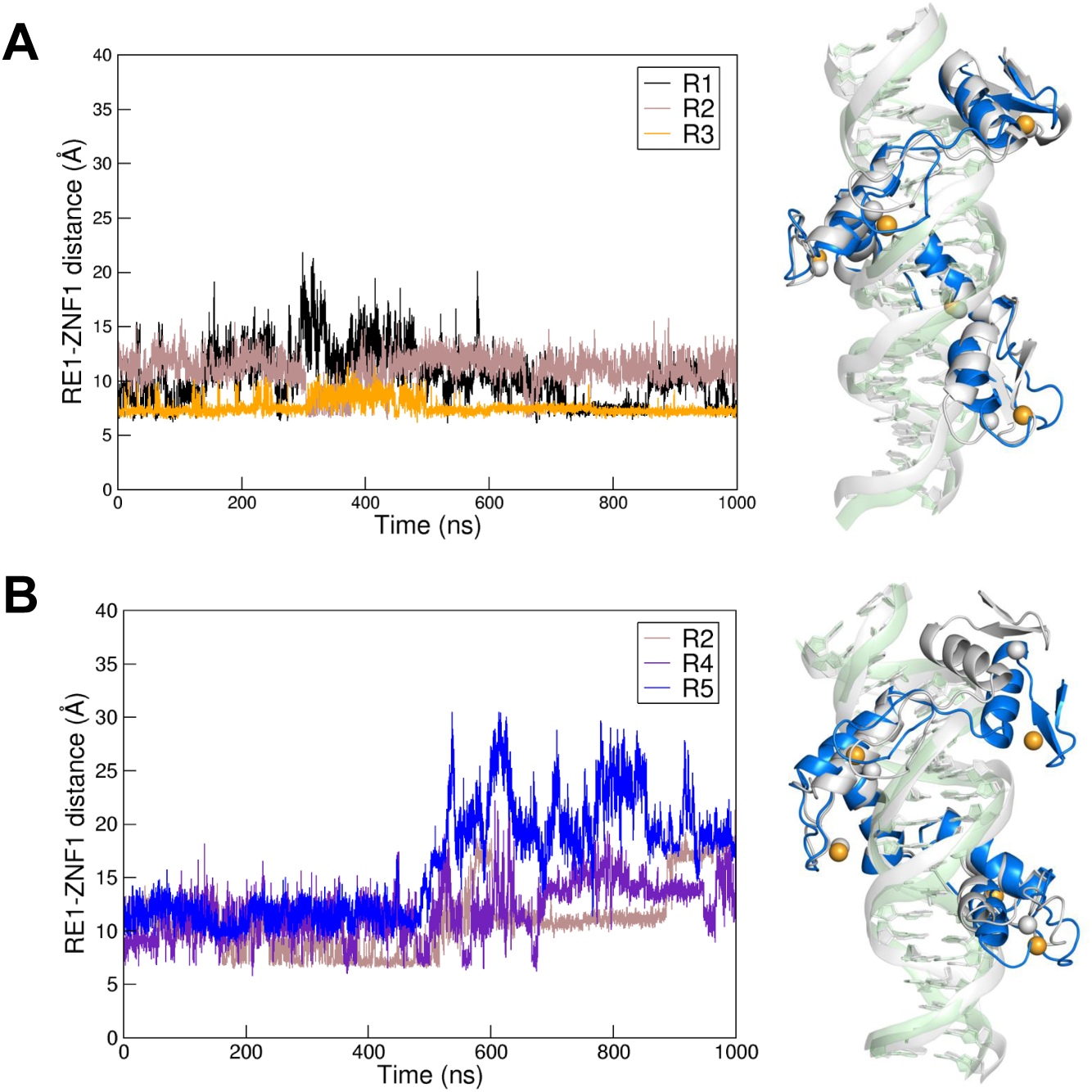
Stability of mREST:RE1 interface in wild-type and S322E phosphomimic complexes. **A**. *Left:* Time evolution of the RE1–ZNF1 distance for three WT replicated simulations. *Right:* Structural superimposition of mREST:RE1 wild-type (WT) complex configurations at the beginning (grey) and at the end (green DNA, blue protein) of one simulated trajectory. **B**. *Left:* RE1–ZNF1 Time evolution of the RE1–ZNF1 distance in three S322E replicas. Right: Structural superimposition of REST:RE1 S322E mutant configurations at the beginning (grey) and at the time point corresponding to the maximum RE1–ZNF1 separation observed in the three trajectories.

To evaluate the potential destabilizing effects of phosphorylation-mimicking mutations, two mREST variants were generated: S322E and T294E, in which S322 and T294, respectively, were substituted with Glutamate. Each mutant was simulated in nine independent 1-µs MD replicas. For the S322E mutant, three replicas (R2, R4, R5) showed an increase in the COM–phosphate distance to 11.2 ± 3.3 Å (R2), 11.8 ± 2.5 Å (R4), and 16.1 Å (R5), with maximum values reaching 24.8 Å, 24.3 Å, and 30.6 Å, respectively, indicating dissociation of the first Zn-finger domain from its DNA binding site (**Figure 7B**). Three other replicas (R7, R8, R9) exhibited structural irregularities, prompted by DNA termini fragilization, leading to complete system collapse in R9 (**Figure S14**). The remaining three trajectories (R1, R3, R6) maintained stable interactions, with average distances of 8.8 ± 1.7 Å, 7.5 ± 0.8 Å, and 9.9 Å ± 1.8 Å, and lower peak excursions (13.6–17.7 Å) (**Figure S13**). For the T294E mutant, two replicas (R1, R4) displayed interface dissociation, with average distances of 11.9 ± 2.2 Å and 10.3 ± 2.6 Å, and maximum distance values of 22.4 Å and 25.0 Å (**Figure S15A**). Two trajectories (R8, R9) showed DNA fragilization, and five replicas (R2, R3, R5, R6, R7) remained structurally stable, with average distances ranging from 7.8 ± 1.0 Å to 11.6 ± 1.6 Å and peak values between 13.0 and 20.9 Å (**Figure S15B**). Taken together, these results show that both phosphomimic substitutions at S322 and T294 introduce measurable destabilization of the mREST:RE1 interaction interface, but that the interface dissociation and structural fragilization are more probable and of higher impact in S322E.

### The CaMKIV/REST interaction underlies the homeostatic effects of CaMKIV on the excitatory synaptic transmission in primary neurons

CaMKIV is known to couple increases in intracellular Ca^2+^ to compensatory changes in excitatory synaptic transmission (Joseph & Turrigiano, 2017). Having identified a mechanistic link between CaMKIV activation and REST activity and half-life, we investigated the interplay between CaMKIV and REST by studying excitatory synaptic transmission in primary cortical neurons. We knocked down CaMKIV by RNA interference in primary cortical neurons derived from conditional REST knockout mice (Rest^GTi^) in which the REST gene was deleted by transduction of Cre recombinase (Cre) or left intact by transduction with a catalytically inactive Cre deletion mutant (ΔCre) and measured miniature excitatory postsynaptic currents (mEPSCs) by patch-clamp in voltage-clamp configuration. Primary neurons in which CaMKIV was knocked down displayed an increase in mEPSC amplitude and no effects on the EPSC frequency, in agreement with previous reports (Joseph & Turrigiano, 2017). On the other hand, REST KO promoted an increase in excitatory strength with augmented mEPSC frequency and no significant effects on mEPSC amplitude (Pegoraro-Bisogni et al., 2018). Interestingly, the concomitant REST deletion virtually abolished the effects of CaMKIV silencing (interaction), indicating that the Ca^2+^-dependent homeostatic effects of CaMKIV are dependent, at least in part, on the presence of REST (**Figure 8**).

**Figure 8.**
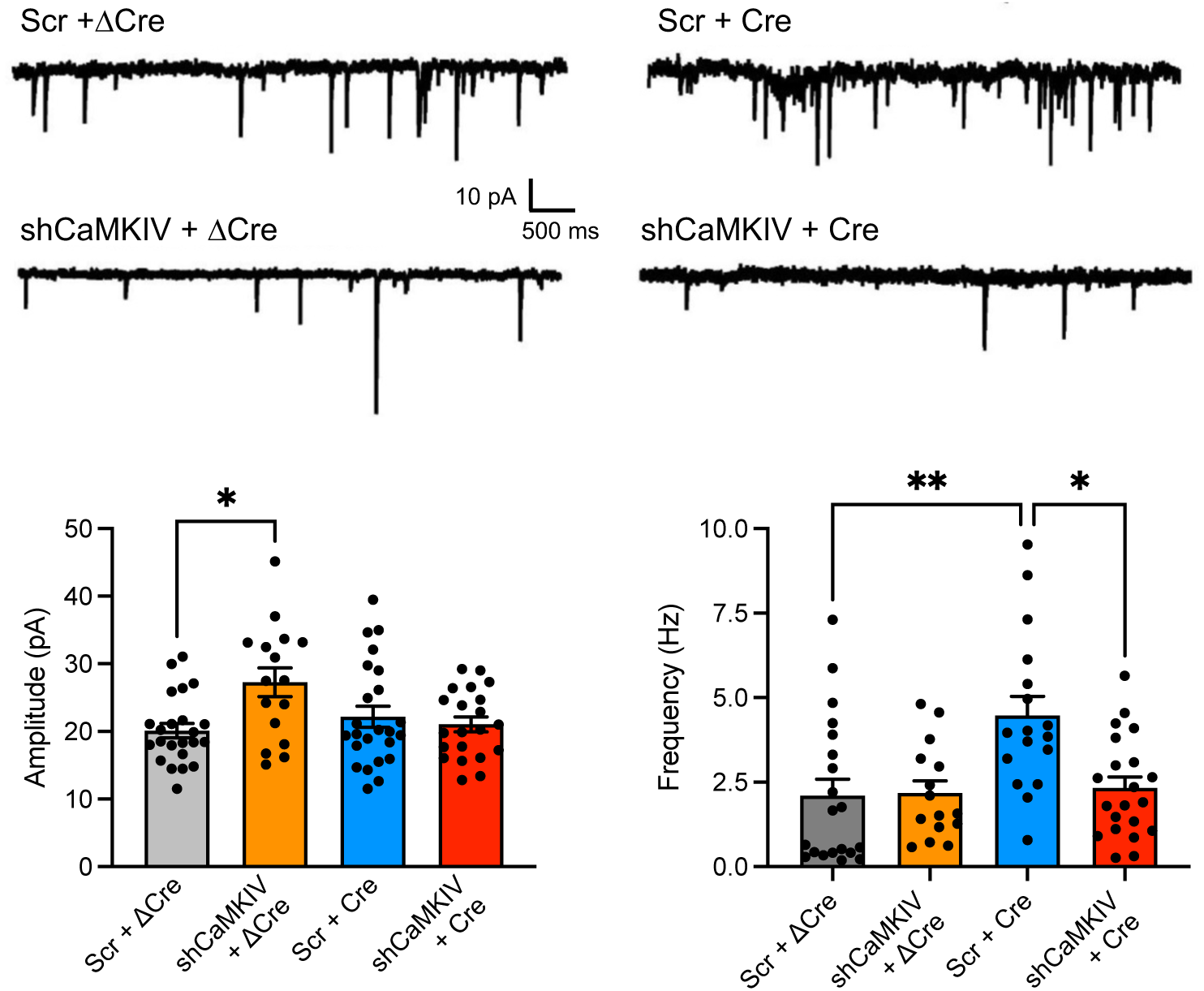
The effects of CaMKIV knockdown on excitatory synaptic transmission in primary cortical neurons are abolished by the conditional knockout of REST/NRSF. **A.** Representative traces of mEPSCs recorded in primary cortical neurons that were knocked out for REST (*Rest*^Cre^) or kept WT (*Rest*^ΔCre^) and co-transduced with either *CaMKIV*^Scr^ or *CaMKIV*^Sh^ expressing lentiviruses. **B.** Mean (± sem with individual experimental points) of the mEPSC amplitude (*left*) and frequency (*right*) in *Rest*^ΔCre^ + *CaMKIV*^Scr^ (n=20), *Rest*^ΔCre^ + *CaMKIV*^Sh^ (n=15), *Rest*^Cre^ + *CaMKIV*^Scr^ (n=17), *Rest*^Cre^ + *CaMKIV*^Sh^ (n=21) neurons from n=3 independent neuronal preparations. Amplitude: *p=0.0172, *CaMKIV*^Scr^ *vs CaMKIV*^Sh^ with *Rest*^ΔCre^; Frequency: **p=0.071, *Rest*^ΔCre^ *vs Rest*^Cre^ with *CaMKIV*^Scr^; *p=0.0126, *CaMKIV*^Scr^ *vs CaMKIV*^Sh^ with *Rest*^Cre^; 2-way ANOVA/Tukey’s tests.

## DISCUSSION

Mature neurons can regulate their network activity to maintain a target level over time through homeostatic plasticity mechanisms regulating intrinsic excitability and synaptic transmission. Several distinct homeostatic transcriptional and posttranslational mechanisms have been identified (Michetti & Benfenati, 2024). They are triggered by a proxy of hyperactivity that most studies indicate as the cytosolic Ca^2+^ concentration, representing an integrated activity signal. Calcium increases activate CaM-dependent enzymes, including the array of CaMKs that, in turn, act at the posttranslational and transcriptional levels, modifying function and expression of ion channels and synaptic proteins (Turrigiano et al., 2011). A key role in the activity-regulated transcriptional mechanisms is played by transcription factors that can be directly stimulated by hyperactivity (e.g., cFos and Arc; Qiu et al., 2022) or activated by phosphorylation. Among CaMKs, CaMKIV, once activated by CaMKK, enters the nucleus and phosphorylates a series of transcription factors such as CREB and MeCP2. A prominent transcription factor that responds rapidly and directly to hyperactivity is REST, an epigenetic master regulator that acts by repressing transcription of a large cluster of neuron-specific genes by interacting with RE1 motifs in their promoters (Ballas & Mandel, 2005; Baldelli & Meldolesi, 2015). In addition to the fundamental role of REST downregulation during neuronal differentiation, which allows the full expression of the neuronal phenotype, REST acts as an important homeostatic regulator in mature neurons that preserves them from hyperactivity and senescence. Dysregulation of REST in terms of overexpression or cell mislocalization was found to associate with brain diseases, such as epilepsy, stroke, neuroinflammation, and neurodegeneration (Buckley et al., 2010; Lu et al., 2014; McClelland et al., 2011, 2014; Hall et al., 2024; Carminati et al., 2021; Buffolo et al., 2021; Natali et al., 2023; Carminati et al., 2020; see Hwang & Zukin, 2018 for review). These considerations emphasize the importance of a tight regulation of REST-DNA interactions, nuclear-cytoplasmic shuttling, and cytoplasmic fate.

REST was reported to be phosphorylated by CK1 and ERK1/2 at distinct sites in its C-terminal region within phosphodegron sequences that direct the protein to the proteasome degradation machinery (Westbrook et al., 2008; Guardavaccaro et al., 2008; Kaneko et al., 2014; Nesti et al., 2014). These mechanisms are thought to be important to decrease cell REST levels in neural progenitors to permit neuronal differentiation. As calcium is believed to be the most likely activator of the homeostatic responses mediated by REST, and REST displays several consensus sequences for CaMKs, we have analyzed the possibility that REST is phosphorylated by CaMKs and that phosphorylation modulates its transcriptional activity and/or cellular fate. Indeed, we found that (i) REST is phosphorylated by CaMKIV at a major phosphorylation site between the 5^th^ and 6^th^ ZNF domains (S322); (ii) phosphorylation by CaMKIV inhibits the transcriptional repressor activity of REST by decreasing its affinity for RE1 motifs; (iii) dissociation from DNA favors shuttling of phospho-REST from the nucleus to the cytoplasm where it undergoes protease-mediated degradation; (iv) the homeostatic effects of CaMKIV require the presence of REST.

Structural studies have shown that the consensus sequences for phosphorylation by CaMKs are, in large part, shared by CaMKI, II, and IV and conform to the following minimal motif: Hyd-X-R-X-X-S*/T*-X-X-X-Hyd (Lee et al., 1994; Dale et al., 1995). While the −3 Arg and −5 hydrophobic residue seem important for all kinases, CaMKI prefers a +4 hydrophobic residue, while CaMKII prefers a +2 acidic residue. The motif for CaMKIV is more degenerate with fewer requirements than the other kinases, supporting the designation of CaMKIV as a multifunctional protein kinase. Of the three putative CaMKIV phosphorylation sites (S177, T294, S322) within the 8 ZNF central cluster region of mouse and human REST, analysis of the location of the basic, hydrophobic and acidic residues in the consensus sequences found that the sequences encompassing T294 and S322 conform better to these requirements, in addition to being the most evolutionary conserved of the three putative sites. Such observation was fully confirmed in the prediction for putative phosphorylation sites by the software Musite and NetPhos3.1, in both the mouse and human REST. Moreover, the REST peptide linkers encompassing the T294 and S322 putative sites share a strong analogy with other DNA-binding transcription factors containing ZNF clusters organized in arrays joined by short 5-7-mer linker sequences. In the 4 ZNF-containing transcription factor Ikaros, for example, ZNF domains are joined by three 5-mer peptides starting with a phosphorylatable residue that, if phosphorylated, impairs the binding of the transcription factor to DNA (Jantz & Berg, 2003). Indeed, MS analysis of in vitro phosphorylated recombinant REST revealed a detectable phosphate incorporation in S322 that thus appears to be the major CaMKIV site in REST.

To explore the mechanistic basis by which phosphorylation modulates REST-DNA binding affinity, we employed atomistic molecular dynamics (MD) simulations of the mREST:RE1 complex in both WT and phosphomimic variants. The simulations revealed that the S322E substitution, mimicking CaMKIV-mediated phosphorylation, induces a notable structural destabilization at the protein-DNA interface. In three independently simulated trajectories over nine, we observed dissociation of the first Zn-finger domain from its DNA binding site, a phenomenon that was totally absent in WT simulations. Furthermore, in several S322E trajectories, destabilization extended beyond the local interface, leading to fragilization of the DNA termini and a collapse of the full complex. Although a comparable loss of interface stability was observed also in some of the T294E simulations, dissociation events occurred less frequently and were not accompanied by complex-wide structural collapse, indicating a more limited effect of the mutation on DNA binding.

These computational findings align with prior experimental observations showing that phosphorylation within or near Zn-finger domains can impair DNA-binding affinity. In particular, phosphorylation of conserved linker sequences in Cys2-His2-type Zn-finger proteins, such as Ikaros and Sp1, has been shown to abolish DNA binding during mitosis by introducing conformational constraints or electrostatic repulsion (Dovat et al., 2002; Jantz & Berg, 2004; Rizkallah et al., 2011). Consistent with this mechanism, our biochemical experiments and MD simulations on mREST mutants demonstrate that a phosphomimic substitution between ZNF5 and ZNF6 within the Zn-finger module directly destabilizes the coordination geometry and its contact with DNA. To our knowledge, this represents the first MD-based evidence of a site-specific post-translational modification within a Zn-finger disrupting the structural stability of a DNA–protein complex at atomic resolution. The simulation results provide structural validation for the biochemical finding that CaMKIV-mediated phosphorylation of REST at Ser322 reduces its affinity for RE1 motifs, facilitating its dissociation from chromatin. Importantly, the absence of collapse or dissociation in WT simulations highlights the specificity of the phosphomimic effect and supports a model in which phosphorylation acts as a trigger for nuclear export and cytosolic degradation. The atomistic-level insight offers a mechanistically distinct layer of control of REST activity directly acting through the DNA-binding domain, in addition to the previously reported regulation via C-terminal phosphodegrons.

Release of CaMKIV-phosphorylated REST from DNA causes its outflow to the cytoplasm, where it undergoes degradation. There are similarities and differences with respect to previous reports on REST phosphorylation. In fact, both CK1 and ERK1/2 are fundamental signaling pathways to achieve REST downregulation during neural development. Their phosphorylation creates phosphodegrons in the C-terminal region of REST, far from the DNA-interacting ZNF domains, that directly target it to the proteasome (Westbrook et al., 2008; Guardavaccaro et al., 2008; Kaneko et al., 2014; Nesti et al., 2014). Given the location of the CaMKIV phosphorylation site within the ZNF domain, REST degradation is likely the consequence of its dissociation from DNA and cytoplasmic migration, rather than the result of the generation of a phosphodegron recruiting ubiquitin ligases. Our results show that CaMKIV-phosphorylated REST is not targeted to the proteasome. Moreover, REST degradation is resistant to the inhibition of organelle acidification by bafilomycin A1, indicating that it does not depend on low pH active proteases present in lysosomes and autophagosomes. Rather, its degradation is blocked by leupeptin, which mostly inhibits lysosomal proteases, which have recently been shown to be active also in extra-lysosomal cytosolic localizations (Conesa-Bakkali et al., 2025). Thus, the available evidence suggests that REST degradation could be mediated by either extra-lysosomal proteases or lysosomal proteases that do not require acidic pH (Curevo et al., 1997).

The interaction between two recognized prominent homeostatic factors, CaMKIV and REST, that are co-activated in response to intracellular Ca^2+^ fluxes during hyperactivity is physiologically of interest. REST responds very rapidly to hyperactivity with increased expression, migration into the nucleus, and repression of target activity genes (Pozzi et al., 2013; Prestigio et al., 2021). CaMKIV is highly expressed in neurons, where it acts downstream of Ca^2+^ signaling. It is activated by the CaMKK phosphorylation, migrates into the nucleus, and modulates transcription of various genes by phosphorylating transcription factors (Xu et al., 2024). The importance of the CaMKIV/REST interplay in the homeostatic effects of CaMKIV is further underlined by the observation that the CaMKIV-induced changes in excitatory synaptic strength (Joseph & Turrigiano, 2017) are virtually abolished by the genetic deletion of REST. As calcium is believed to be the most likely activator of the homeostatic responses mediated by both REST and CaMKIV, the results suggest that the homeostatic responses of neurons to hyperactivity may require an interplay between CaMKIV and REST, which may define the temporal window of activation of the homeostatic response before the return to basal conditions.

The timing of epigenetic changes and transcriptional effects induced by REST is a key issue, since transcriptional effects can last for long times and thus require a negative regulation of REST activation and overexpression to return to baseline. Termination of REST activity may consist of a sequence of events started by phosphorylation by CaMKIV in the nucleus, dissociation of phospho-REST from the DNA, shuttling to the cytoplasm, followed by degradation. Thus, the elucidated CaMKIV-REST interplay can play an important regulatory role, bringing back the homeostatically increased REST levels to a physiological transcriptional profile after the inhibition of hyperactivity. Such a mechanistic model, shown in **Figure 9**, defines two phases of the homeostatic response to hyperactivity, namely an early phase triggering long-lasting transcriptional effects to decrease neuronal activity, and a subsequent phase in which the transcriptional activators/repressors are switched off.

**Figure 9.**
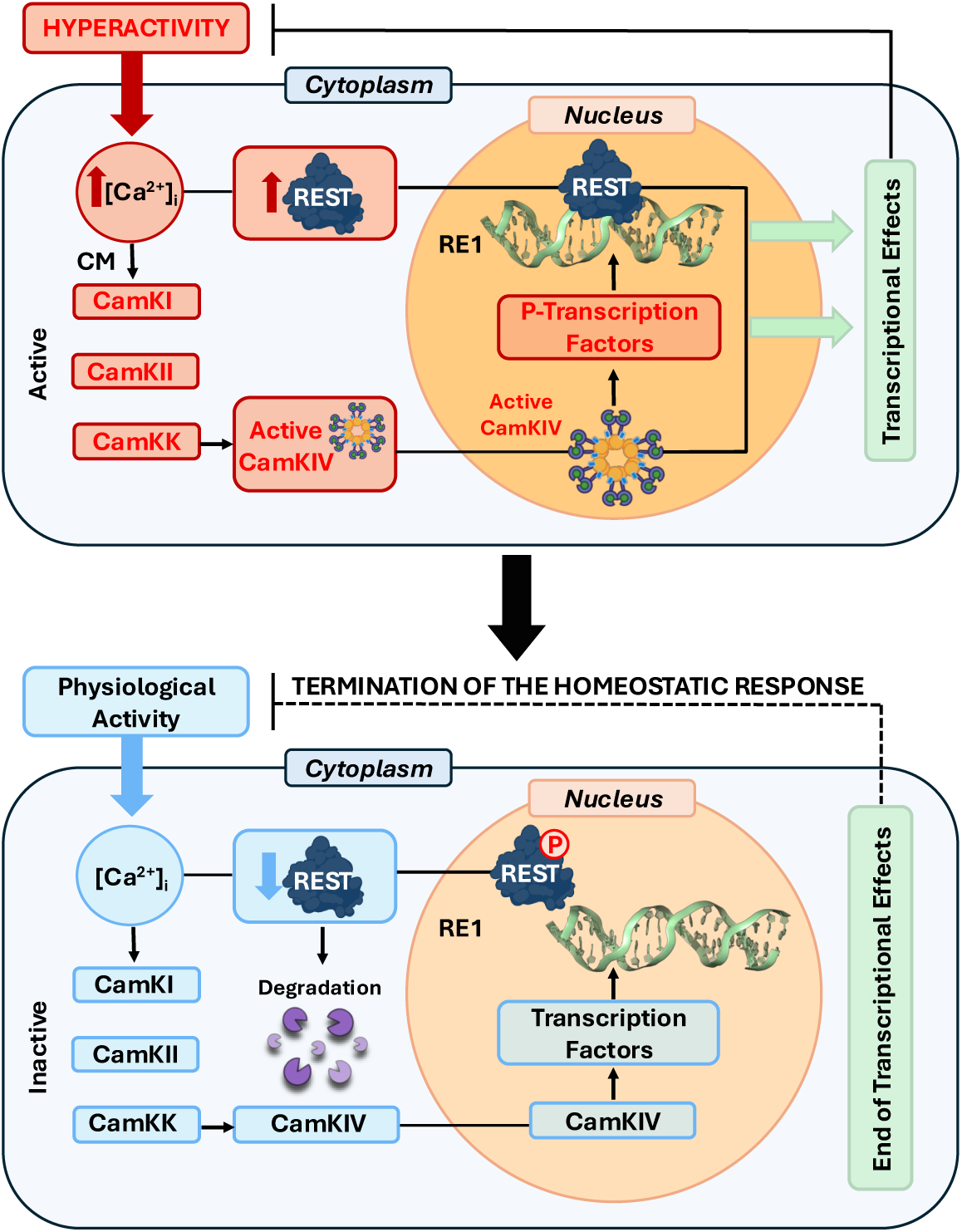
Proposed model for the effects of REST phosphorylation by CaMKIV. Calcium entry during hyperactivity activates both REST expression and CaMKIV. REST homeostatically downregulates intrinsic excitability and excitatory synaptic transmission by acting at the transcriptional level in the nucleus. Meanwhile, CaMKIV, activated by Ca^2+^/calmodulin, translocates to the nucleus and phosphorylates REST at S^322^ adjacent to the 5^th^ Zn-finger domain, which participates in the interaction of the transcription factor with DNA. Phosphorylated REST has a reduced affinity for the RE1 sequences present in the promoter region of its target genes, and, upon unbinding, it is shuttled back to the cytosol, where it is eventually targeted for degradation. This way, the homeostatic plasticity response triggered by hyperactivity will terminate once activity has been normalized and the increased REST levels have returned to baseline.

In conclusion, we found a control mechanism that may contribute to closing the homeostatic transcriptional loop in response to hyperactivity and that involves two well-known transcriptional regulators. A deeper understanding of REST phosphorylation dynamics by CaMKIV could uncover novel regulatory nodes with therapeutic potential in neurological disorders.

## Acknowledgements

We are grateful to Drs. Gail Mandel (Vollum Institute, Portland, OR), Thomas Floss (Helmholtz Zentrum München, Deutsches Forschungszentrum für Gesundheit und Umwelt, Neuherberg, Germany), and the German Gene Trap Consortium (GGTC-Partners) for providing us with the REST conditional knockout mice. We are also grateful to Drs Angus Nairn (Yale University School of Medicine, New Haven, MA) and Taku Kaitsuka (Kumamoto University, Kumamoto, Japan) for kindly providing the DNA constructs for WT, catalytically active, and kinase-dead CaMKs. We acknowledge ISCRA for awarding this project access to the LEONARDO supercomputer, owned by the EuroHPC Joint Undertaking, hosted by CINECA (Bologna, Italy). We also gratefully acknowledge the Data Science and Computation Facility and its Support Team for their support and assistance on the IIT High Performance Computing Infrastructure (Genova, Italy). We also thank: Drs. Maria Sabina Cerullo and Alessandra Romei for help in some initial experiments; Dr. Pietro Baldelli for critical reading of the manuscript; drs. Riccardo Navone (Istituto Italiano di Tecnologia, Genova, Italy), Laura Emionite, and Michele Cilli (IRCCS Ospedale Policlinico San Martino, Genova, Italy) for assistance in breeding the mice; Drs. Diego Moruzzo and Arta Mehilli (Istituto Italiano di Tecnologia, Genova, Italy) for help in genotyping; Drs. Ilaria Dallorto and Rossana Ciancio (Istituto Italiano di Tecnologia, Genova, Italy) for administrative assistance.

## Funding

The study was supported by research grants from the Italian Ministry of Health (RF-2021-12374404, GR-2019-12370176, and PNRR-MR1-2022-12376528), Compagnia di San Paolo Torino (Trapezio #118787 and #124077), the Italian Ministry of University and Research (PRIN 2020-WMSNBL and PRIN 2022-XWEYBN), and by IRCCS Ospedale Policlinico San Martino (Ricerca Corrente) and #NEXTGENERATIONEU (NGEU) National Recovery and Resilience Plan (NRRP), project MNESYS (PE0000006) – A Multiscale integrated approach to the study of the nervous system in health and disease (DN. 1553 11.10.2022 to HTS and AF) and project B83C22004960002 Light4Brain (to FB).

## Author Contributions

HTS, AR, DS and EC run molecular biology, biochemistry, viral vector preparation, and immunohistochemical analyses; BC and LM setup the REST-DNA model, run the MD simulations, and analyzed the data; AA, NL and ADF carried out the mass spectrometry analysis; HTS and AF carried out the REST degradation experiments; MC carried out in vitro electrophysiological experiments and analysis; FB, AR and AF conceived the study, supervised the experiments, and analyzed the experimental data; FB wrote the manuscript and supported the research; all authors critically discussed the experimental results and contributed to manuscript writing and revision.

## MATERIALS AND METHODS

### Materials and experimental animals

#### Experimental animals

Wild-type C57BL/6J male mice were purchased from Charles River (Calco, Italy). GTinvRest (RestGTi) mice (Nechiporuk et al. 2016), obtained by Gail Mandel (Portland, OR) and the German Gene Trap Consortium (GGTC-Partners), were kept on a C57BL/6J background in homozygosity under conditions of environmental enrichment. Mice were maintained on a 12:12 h light/dark cycle at constant temperature (22 ± 1 °C) and humidity (60 ± 10%), with water and pellet diet *ad libitum.* All the experiments were strictly following the guidelines of the European Community Council (Directive 2014/63/EU of May 15, 2014) and were approved by the Italian Ministry of Health (Authorization #427/2021-PR).

#### Materials

W7 (N-(6-aminohexyl)-5-chloro-1-naphthalenesulfonamide hydrochloride), KN92, KN93, bicuculline, CGP58845, D-APV, and tetrodotoxin were from Tocris (Bristol, UK). Cyclosporin A, cycloheximide, MG132, leupeptin, and bafilomycin A1 were from Merck Life Science (Milan, Italy). All other chemicals, unless otherwise specified, were from Sigma-Aldrich (Milano, Italy)

### Primary neuron cultures

Cultures of primary cortical neurons were obtained either from wild-type C57Bl/6 or GTinvRest mice embryos, as previously described (Baldelli et al., 2007; Chiappalone et al., 2009; Valente et al., 2016). These cultures are very homogeneous, contain ∼75% excitatory neurons with a recognizable pyramidal shape, ∼20-25% glutamic acid decarboxylase (GAD)-positive inhibitory neurons, very low percentages of astrocytes (5-7%), and no microglial cells due to the embryonic nature of the preparation. In brief, cerebral cortices from E17–E18 embryos were dissociated by enzymatic digestion in 0.125% trypsin for 20 min at 37 °C and then triturated with a fire-polished Pasteur pipette. Primary cultures of dissociated cortical neurons were subsequently plated onto poly-L-lysine (0.1 mg/mL, Merck Life Science)-coated 18-mm glass coverslips (6 × 10^4^ cells/coverslip for immunofluorescence and electrophysiology experiments) and poly-L-lysine (0.01 mg/mL)-coated 35-mm wells (5 × 10^5^ cells/well for molecular biological and biochemical experiments). Neurons were maintained in a culture medium consisting of Neurobasal, supplemented with B27 (1:50 v/v), Glutamax (1% w/v), penicillin-streptomycin (1%) (ThermoFisher Scientific, Monza, Italy), and kept at 37 °C in a 5% CO_2_ humidified atmosphere for 14 days before harvesting.

### Manipulations of CaM-dependent pathways in primary cortical neurons

Neurons were treated for 24 h prior to harvesting with the following compounds: N-(6-aminohexyl)-5-chloro-1-naphthalenesulfonamide hydrochloride (W7; 20 µM), Cyclosporin A (CsA; 1 µM), N-[2-[N-(4-chlorocinnamyl)-N-methylaminomethyl]phenyl]-N-(2-hydroxyethyl)-4-methoxybenzene sulfonamide (KN93; 10 µM), and 2-[N-(4-methoxybenzenesulfonyl)]amino-N-(4-chlorocinnamyl)-N-methylbenzylamine phosphate (KN92; 10 µM). All drugs were obtained from Tocris. Dimethyl sulfoxide (DMSO; Cat. No. D2650, Sigma-Aldrich, Milan, Italy) was used as a vehicle control according to the experimental conditions. To specifically alter their CaMK activities, HEK293T cells were alternatively transfected with the following plasmids using Lipofectamine 2000 (ThermoFisher Scientific):

pCMV-HA CaMK1 (WT, CA or KD; kind gift of Dr Taku Kaitsuka, Kumamoto University, Kumamoto, Japan);

pBS-EGFP-MT-CaMK2A (EMBL Database #AJ890283), pBS-EGFP-MT-CaMK2A K42R (CA), and pBS-EGFP-MT-CaMK2A T286D (KD), kind gifts from Dr. Agnes Thalhammer (Istituto Italiano di Tecnologia, Genova, Italy);

pRSV-CaMK4WT (#45062; Addgene, Watertown, MA), pRSV-CaMK4 CA (1-313) (#45063; Addgene, Watertown, MA), and pRSV-CaMK4 KD was mutagenized using pRSV-CaMKIV WT as template.

### REST expression in Hek293 cells

HEK293T cells at approximately 70% confluency in 15-cm plates were transfected with 9.2 µg of pCS-myc-mREST per plate (Pozzi et al. 2013). Cells were washed and collected 48 h after transfection in phosphate-buffered saline (PBS). Cell lysates were prepared using the Pierce c-Myc-Tag Magnetic IP/Co-IP Kit (ThermoFisher Scientific) according to the manufacturer’s instructions. Briefly, cells were collected by centrifugation at 100xg for 5 min. Pellets were resuspended in 6 mL of Co-IP buffer (ThermoFisher Scientific) and incubated on ice for 10 min before centrifugation at 18,500xg for 15 min at 4 °C, and the supernatant was sonicated at 25% power for 5 s on ice. The protein concentration was determined by the bicinchoninic assay using Pierce™ BCA Protein Assay Kit (ThermoFisher Scientific). The remaining total lysates were concentrated using Amicon Ultra Centrifugal Filter (50 kDa MWCO; Merck Life Science) at 18,500xg for 20 min at 4 °C. Anti-c-Myc magnetic beads were washed three times in Tris-buffered saline containing Tween-20 (TBS-T) (25 mM Tris, 0.15 M NaCl, 0.05% v/v Tween-20).

### On-bead REST phosphorylation by CaMKIV

Approximately 200 µg of the concentrated total lysate was incubated with Pierce^TM^ anti-c-Myc magnetic beads (ThermoFisher Scientific) that were pre-washed according to the manufacturer’s instructions, diluted 1:15 with TBS-T buffer, and incubated at room temperature (RT) for 30 min under rotation. Beads, collected on the magnetic stand and washed in TBS-T three times, were resuspended in alkaline phosphatase buffer (5 mM Tris-Cl pH 8.7, 10 mM NaCl, 1 mM MgCl_2_, 0.1 mM DTT) supplemented with protease inhibitor cocktail (Merck Life Science) and treated with 25 U of alkaline phosphatase (Merck Life Science) for 30 min at 30 °C. The reaction was stopped by removing the supernatant on the magnetic stand and washing the beads twice with 200 µL of TBS-T buffer supplemented with 10 mM EDTA and protease inhibitors. The beads were subsequently resuspended in *in vitro* phosphorylation assay buffer (50 mM HEPES at pH7.4, 10 mM MgCl_2_, 1 mM EGTA, 0.4 mM DTT), supplemented with protease and phosphatase inhibitors, 60 µM Mg^2+^-ATP, and 600 nM calmodulin (Merck Life Science). Either 1 µg of active recombinant CamKIV (Merck Life Science) or vehicle (negative control) was added to the samples, and the mix was incubated for 1 h at RT under rotation. The on-bead kinase assay was stopped by removing the supernatant and washing the beads with an ice-cold assay buffer (10 mM Tris-HCl at pH 7.4, 75 mM NaCl, 20 mM EDTA) supplemented with protease and phosphatase inhibitors. The bound fractions were eluted in 2 x Laemmli buffer (4% SDS, 20% glycerol, 120 mM Tris-HCl pH 6.8, 200 mM DTT).

### Protein Digestion and Mass Spectrometry

The protein content resulting from the on-bead phosphorylation assay was reduced with DTT, alkylated with iodoacetamide, then digested overnight with trypsin following a well-established protocol (Castagnola et al., 2024; Liessi et al., 2021). The proteomic analysis was conducted on a Thermo Exploris 480 orbitrap system coupled with a Dionex Ultimate 3000 nano-LC system. After trapping and desalting, the tryptic peptides were loaded on an Aurora C18 column (75 mm x 250 mm, 1.6 µm particle size) nanocolumn (Ion Opticks, Fitzroy, Australia) and separated using a linear gradient of acetonitrile in water (both added with 0.1% formic acid), from 3% to 41% in 1 h, followed by column cleaning and reconditioning. The flow rate was set to 300 nL/min. Total run time was 1.5 h. Injection volume was set to 1 µL. Peptides were analyzed in positive ESI mode, using a capillary voltage set to 2.0 kV. The RF lens was set to 40%, and the AGC target was set to 300%. Data acquisition was performed in Data Dependent mode (DDA) MS/MS spectra were acquired in HCD mode. All the collected MS/MS spectra were analyzed using Proteome Discoverer software. Cysteine carbamido-methylation was selected as a fixed modification; acetylation of protein N-Term, methionine oxidation, and phosphorylation at serine, threonine, and tyrosine were selected as variable modifications. Positive protein identifications were retained at a 1% false discovery rate (FDR) threshold and with at least two peptides matching the protein sequence. Unless otherwise specified, all reagents were purchased from Sigma Aldrich, and LC-MS solvents from ThermoFisher. All Instruments and software were purchased from ThermoFisher Scientific.

### Inhibition of REST degradation pathways

HEK293T cells at approximately 70% confluency in 3.5-cm plates were transfected with 0.5 µg of either pRSV-CaMKIV (full-length, WT) or pRSV-CaMKIV (1-313; constitutively active) using Lipofectamine 2000 (ThermoFisher Scientific) and DMEM (ThermoFisher Scientific), supplemented with FBS (10%; ThermoFisher Scientific) and penicillin-streptomycin (1%; ThermoFisher Scientific), and were kept at 37 °C in a 5% CO_2_ humidified atmosphere. One day after transfection, the medium was replaced with DMEM (ThermoFisher Scientific) with cycloheximide (35 µM; Merck Life Science), 5% FBS, and 1% penicillin-streptomycin. Cells were kept at 37 °C in a 5% CO_2_ humidified atmosphere. After 3 h, cells were washed three times in DMEM/5% FBS/1% penicillin-streptomycin and incubated in the same medium containing vehicle (0.1 % DMSO; Merck Life Science), MG132 (5 µM; Merck Life Science), Bafilomycin A1 (10-100 nM; Merck Life Science), or Leupeptin (40 µM; Merck Life Science). Following 16 h of incubation at 37 °C in a 5% CO_2_ humidified atmosphere, cells were washed three times in PBS and processed as described for Western blotting.

### Site-directed mutagenesis of REST phosphorylation sites

Nucleotides encoding Threonine-294 and Serine-322 were mutated to either alanine or glutamic acid using the protocol of DpnI-mediated site-directed mutagenesis by Fisher and Pei (1997). PCR amplification was performed on the template plasmid containing wild-type mREST cDNA in pEZ-Lv229-mREST-IRES2-mCherry constructs with the following protocol: 95 °C for 30 s, 18 cycles of: 95 °C for 30 s, 55 °C for 1 min, 68 °C for 20 min using the pair of primers for each mutated site. The methylated template DNA contained in the PCR products was digested by incubation with 10 U of DpnI for 2 h at 37 °C. A fraction (1/50) of the digested PCR product was used to transform TOP10-competent cells by heat shock at 42 °C for 30 s, incubated in SOC medium for 30 min at 37 °C, and then plated on an LB-agar plate with ampicillin (100 µg/mL). Mini preps from 4 colonies were prepared for full-length sequencing to identify the correct clone with mutated sites.

### Western blotting

Primary cortical neurons or HEK293T cells were used to obtain total cell lysates. Cells were collected in Lysis buffer (150 mM NaCl, 1% NP-40, 0.5% sodium deoxycholate, 0.1% SDS, 2 mM EDTA, 50 mM Tris-HCl pH 7.4) enriched with protease and phosphatase inhibitor cocktails (Merck Life Science, Milan, Italy). Following a 10-min incubation on ice, cell lysates were centrifuged at 18,500xg for 15 min at 4 °C. The supernatant was collected and sonicated at 25% power for 5 s on ice. The BCA assay (ThermoFisher Scientific) was used to determine protein concentration. Equivalent protein amounts were subjected to SDS-PAGE on 7.5% polyacrylamide gels and blotted onto nitrocellulose membranes (Whatman, Maidstone, UK). Blotted membranes were blocked for 1 h using 1% BSA (Merck Life Science) in TBS (10 mM Tris, 150 mM NaCl, pH 8.0) together with 0.1% Triton X-100, and incubated overnight at 4 °C with the following primary antibodies: anti-CaMKI (#ab68234 Abcam, Cambridge, UK), anti-CaMKII (#07-1496, Merck Life Science), anti-CaMKIV (#38 610275 BD Biosciences, Milano, Italy), anti-REST (#07-579, Merck Life Science), anti-GAPDH (#2118, Cell Signalling/Euroclone, Milano, Italy), anti-Calnexin (#ADI-SPA-860, Enzo Lifesciences, Farmingdale, NY). Membranes were washed and incubated at RT for 1 h with the following peroxidase-conjugated secondary antibodies: goat anti-mouse IgG (1:10,000; #32430; ThermoFisher Scientific) and goat anti-rabbit IgG (1:5000; #32460; ThermoFisher Scientific). Bands were revealed using the ECL chemiluminescence detection system (ThermoFisher Scientific), acquired on the iBright Imaging System (ThermoFisher Scientific), and the immunoreactivity quantification was performed by densitometric analysis with ImageJ.

### Real-time qPCR

Total cellular RNA was extracted with TRIzol (ThermoFisher Scientific), and RNA concentration was quantified using the Nanodrop-1000 spectrophotometer (ThermoFisher Scientific). cDNA was synthesized starting from 0.25 µg RNA with SuperScript IV Reverse Transcriptase kit (ThermoFisher Scientific) according to the manufacturer’s instructions. Gene expression was measured by real-time qPCR using SsoAdvanced Universal SYBR Green Supermix (Bio-Rad Laboratories, Milano, Italy) on a C1000 Touch Thermal Cycler (Bio-Rad) on a CFX96 Real-Time System following the manufacturer’s protocol, with the following protocol: 5 min at 95 °C; 10 s at 95 °C; 20 s at 60 °C; 10 s at 72 °C for 45 cycles; melting curve (heating ramp from 55 to 95 °C) to check for amplification specificity. The following primers (final concentration 0.25 µM) were used:

REST_F: 5’- GAACCACCTCCCAGTATG −3’

REST_R: 5’- CTTCTGACAATCCTCCATAG −3’

CaMKI_F: 5’- ATCAAGGAAGTCAGGGTTT −3’

CaMKI_R: 5’- GCAGTGAAGAGTGAGAGG −3’

CaMKII_F: 5’- ACTTCCTTCCACCACTTC −3’

CaMKII_R: 5’- TGAGATACAGCATTCCATACA −3’

CaMKIV_F: 5’- CAGTTCATGTTCAGGAGAAT- 3’

CaMKIV_R: 5’- AATGTAGTCAGCCGTTTC −3’

Gapdh_F: 5’- AGGTCGGTGTGAACGGATTTG −3’

Gapdh_R: 5’- TGTAGACCATGTAGTTGAGGTCA −3’

The primers used for the mutagenesis were the following:

mRest_T294E_F: 5’-AGCACGTGCGAACTCACGAAGGAGAACGCCCGTAT-3’

mRest_T294E_R: 5’-ATACGGGCGTTCTCCTTCGTGAGTTCGCACGTGCT-3’

mRest_S322E F: 5’-GACACATGCGGACTCATGAAGGTGAGAAGCCAT-3’

mRest_S322E R: 5’-ATGGCTTCTCACCTTCATGAGTCCGCATGTGTC-3’

Relative gene expression was determined using the 2-ΔΔCT method, normalizing data on the *Gapdh* housekeeping transcript (Vandesompele et al., 2002).

### Cell transfection and immunocytochemistry

HEK293T cells on 18-mm coverslips at 70% confluency were transfected with 0.5 µg of pEZ-Lv229-mREST-IRES2-mCherry WT (GeneCopoeia, Rockville MD) or mutagenized constructs therefrom using Lipofectamine 2000 (ThermoFisher Scientific). One day after transfection, cells were fixed in 4% paraformaldehyde in 0.1 M phosphate buffer pH 7.4 (PB) for 20 min at RT. Fixed cells were stained with anti-REST antibody (#MABE1982, clone 12C11) and Hoechst to visualize the nuclear area. Briefly, cells were washed in PBS three times, permeabilized and blocked, and incubated with the primary antibody overnight at 4 °C in 0.1% Triton X-100 and 5% normal goat serum (NGS) in PBS. After four washes in high-salt PBS (20 mM Na_2_HPO_4_/ 20 mM NaH_2_PO_4_/ 0.5 M NaCl) for a total of 20 min, neurons were incubated in the same buffer with Alexa-conjugated secondary antibodies (1:500, ThermoFisher Scientific) and counterstained with Hoechst33342 for nuclear detection. Following a wash in high-salt PBS for 20 min, coverslips were washed once in PBS, once in double-distilled water, and mounted using a Moviol mounting medium. Tilescan images of transfected HEK293T cells were acquired using a 60x objective with a Leica SP8 confocal microscope (Leica Microsystem, Wetzlar, Germany). Colocalization of anti-REST immunostaining signal with Hoechst or with mCherry was analyzed in binarized tilescan images using the JACoP plugin for the FIJI distribution of ImageJ. Colocalization was computed between all channels, with special focus on nuclear localization of anti-REST fluorescence. Manders’ overlap coefficients were extracted as measures of reciprocal channel colocalization alongside Pearson’s correlation coefficient with Costes’ randomization (200 iterations) to test for overlap quality and significance.

### Luciferase assay

HEK293T cells at 50% confluency in a 6-well plate were co-transfected with RE1X3-pGL3 promoter vector (Paonessa et al., 2016), with mutagenized mREST in pEZ-Lv229-mREST-IRES2-mCherry, or with pRSV-CaMKIV(1-313) (#45063, Addgene, Watertown, MA) and the pRL TKSV40 vector expressing *Renilla* luciferase (Promega, Milano, Italy), using Lipofectamine 2000 (ThermoFisher Scientific). Two days after transfection, cells were washed in PBS twice and collected in 1x Passive Lysis Buffer (PLB) from Dual-Luciferase Assay Kit (Promega), supplemented with protease inhibitor cocktail. The lysates were subjected to one free-thaw cycle and were centrifuged at 100xg at 4 °C for 5 min to remove residual cell debris. The supernatants were collected, diluted 1:5 in 1xPLB, and added to multi-wells containing the luciferase substrate LARII. After a first basal reading of luminescence, a second reading was performed after the addition of Stop & Glo Reagent (Promega). All readings of luminescence were made using a Spark Multimodal Microplate Reader (Tecan, Zurich, Switzerland).

### Transduction of primary neurons

#### Generation of lentiviral vectors

VSV-pseudo-typed third-generation lentiviruses were obtained as previously described (De Palma and Naldini, 2002). Briefly, lentiviruses were produced by transient four-plasmid co-transfection into HEK293T cells using the Ca^2+^ phosphate transfection method. Supernatants were collected, passed through a 0.45-μm filter, and purified by ultracentrifugation (75,000xg for 2 h, at 4 °C). Viral vectors were titrated at concentrations ranging from 1×10^8^ to 1×10^9^ transducing units (TU)/mL and used at a multiplicity of infection (MOI) of 5-10.

#### REST RNA interference

For specific knockdown of CaMKs in neurons, short hairpin RNA (shRNA) was cloned by inserting the annealed oligos of shRNA for either CaMKI or CaMKIV into the pLenti-U6-shRNA-PGK-GFP vector between the XhoI and HpaI sites.

The shRNA oligonucleotides used were as follows:

shCaMKIa Sense oligo III: 5’- TGCATTGTAGCCCTGGATGACTTCAAGAGA GTCATCCAGGGCTACAATGCTTTTTTC - 3’

shCaMKIa Antisense oligo III: 5’- CTCGAGAAAAAAGCATTGTAGCCCTGGATGA CTCTCTTGAAGTCATCCAGGGCTACAATGCA - 3’

shCaMKIa Sense oligo IV: 5’- TGGATCAAGCACCCCAACATTTTCAAGAGA AATGTTGGGGTGCTTGATCCTTTTTTC - 3’

shCaMKIa Antisense oligo IV: 5’- CTCGAGAAAAAAGGATCAAGCACCCCAACATT TCTCTTGAAAATGTTGGGGTGCTTGATCCA - 3’

shCaMKIV Sense oligo II: 5’- TGAGAGAATCTTCTTTATGCATTCAAGAGA TGCATAAAGAAGATTCTCTCTTTTTTC - 3’

shCaMKIV Antisense oligo II: 5’- CTCGAGAAAAAAGAGAGAATCTTCTTTATGCA TCTCTTGAATGCATAAAGAAGATTCTCTCA - 3’

shCaMKIV Sense oligo III: 5’- TGGTGTTAAAGAAAACAGTGGTTCAAGAGA CCACTGTTTTCTTTAACACCTTTTTTC - 3’

shCaMKIV Antisense oligo III: 5’- CTCGAGAAAAAAGGTGTTAAAGAAAACAGTGG TCTCTTGAACCACTGTTTTCTTTAACACCA - 3’

scramble Sense oligo: 5’- TGAGAGAATCTTCTTTATGCATTCAAGAGA TTGGGTTGAAGGTGGATCCCTTTTTTC - 3’

scramble Antisense oligo: 5’- CTCGAGAAAAAAGGGATCCACCTTCAACCCAA TCTCTTGAATGCATAAAGAAGATTCTCTCA - 3

#### Transduction of primary neurons

For silencing CaMKs in WT primary cortical neurons, neurons were transduced with lentiviral vectors encoding the respective shRNA together with the GFP reporter (see above). For silencing CaMKIV and/or knocking out REST, primary cortical neurons from RestGTi cKO mice were alternatively transduced with sequences containing active or inactive Cre recombinase (Cre and its deletion mutant ΔCre) that were cloned into pGFP-Lenti using the PGK ubiquitous promoter (Kaeser et al., 2011) together with either scrambled or CaMKIV shRNA using the following vector: pLenti-U6-(scramble/shCaMKIV)-PGK-(ΔCRE/CRE-T2A-GFP). For all the experiments, cKO cortical neurons at 7 days in vitro (DIV) were infected at a multiplicity of infection (MOI) of 10. After 24 h, half of the medium was replaced with fresh medium. Transduction efficiency was. The efficiency of infection was estimated to range between 70% and 90% by counting neurons expressing GFP with respect to the total number of DAPI-stained cells. Primary cortical neurons were infected at 7 DIV. After 24 h, half of the medium was replaced with fresh medium. Experiments were performed 7 days after infection.

### Structural Modeling of the REST:RE1 Complex

#### Atomistic Molecular Dynamics simulations of the wild-type REST-RE1 complex

Structures of the WT REST–RE1 assembly were generated as described in the Appendix (Section *Molecular Modeling of Wild-Type REST* and following). The solution builder plug-in of the CHARMM-GUI web server (Lee et al., 2016, 2019) was utilized for setting up the system and for generating input files to be used in all-atom molecular dynamics (MD) simulations. The DNA-protein assembly was inserted into a rectangular box of TIP3 water molecules (Jorgensen et al.,1983) with a minimum solute-to-box-edge distance of 15 Å, applied along all directions. The total system charge was neutralized by adding Na^+^ and Cl^-^ ions reaching a physiological concentration of 0.15 M. The whole system amounts to ∽112000 atoms. Simulations were performed with NAMD 2.14 (Phillips et al., 2020), using the Amber Ff14SB (Maier et al., 2015) force field for the protein and the BSC1 (Ivani et al., 2016) force field for the nucleic acids. Tetragonal periodic boundary conditions (PBC; de Leeuw et al., 1980) were applied. The Particle Mesh Ewald method (Darden et al., 1993) was used to treat electrostatic interactions and to avoid box surface effects with a maximum grid spacing of 1 Å and sixth-order beta splines. A cutoff of 12 Å was employed for Lennard-Jones interactions, which were smoothly switched off starting at 10 Å. Chemical bond distances involving hydrogen atoms were constrained using the SHAKE algorithm (Ryckaert et al., 1977). A time step of 2 fs was utilized. Before entering the production stage, the system was energy minimized and equilibrated in the isobaric-isothermal ensemble (NPT) with P=1 atm and T=310 K. Pressure and temperature were maintained constant using the Nose-Hoover Langevin piston (Feller et al., 1995) and the Langevin thermostat (Leimkuhler et al., 2009), respectively. Afterwards, three independent, 1-µs-long unbiased all-atom MD simulations were performed in the NPT ensemble. *Clustering of wild-type REST-RE1.* To generate phosphomimic mutant conformations, we extracted snapshots from the equilibrium simulations of WT REST-RE1. Trajectories were individually subjected to clustering analysis employing TTClust4.10.3 (Tubiana et al., 2018). MD frames were aligned using the backbone of both the protein and the DNA as a reference. Pairwise RMSDs were assembled into a matrix and used for hierarchical clustering with the Ward algorithm (Murtagh & Legendre, 2014) to build the linkage matrix, minimizing the increase in within-cluster variance. Structures of cluster centers were visually inspected using the PyMOL Molecular Graphics System (2002, Version 2.0, Schrödinger, LLC, New York, NY), and, for each replica, the center of the most populated cluster was taken as the representative conformation of that MD run.

#### Atomistic Molecular Dynamics Simulations of REST phosphomimics

Two distinct phosphomimic mutants were generated by varying the amino acid identity of positions T154 and S192 (T294 and S322 in the experimental system, respectively) into glutamic acid. All simulation procedures and parameters were the same as for the WT system. A total of 18 independent, 1 µs-long, unbiased all-atom MD simulations, 9 per mutated system, were performed in the NPT ensemble.

### Patch-clamp electrophysiology

Whole patch-clamp recordings were made from primary cortical neurons as previously described (REF) using a Multiclamp 700B/Digidata 1400A system (Molecular Devices, Sunnyvale, CA). Patch pipettes, prepared from borosilicate glass, were pulled and fire-polished to a final resistance of 4-5 MΩ when filled with a standard internal solution. For all the experiments, neurons were maintained in standard extracellular Tyrode solution containing (in mM) 140 NaCl, 2 CaCl_2_, 1 MgCl_2_, 4 KCl, 10 glucose, and 10 HEPES (pH 7.3 with NaOH). For recording miniature excitatory postsynaptic currents (mEPSCs), bicuculline (30 µM), CGP58845 (10 µM), D-APV (50 µM), and tetrodotoxin (TTX; 300 nM) were added to the extracellular solution to block GABA_A_, GABA_B_, NMDA receptors and generation and propagation of spontaneous APs. The internal solution (K-gluconate) used for recording mEPSCs in voltage-clamp configuration contained (in mM): 126 K gluconate, 4 NaCl, 1 MgSO_4_, 0.02 CaCl_2_, 0.1 BAPTA, 15 glucose, 5 HEPES, 3 ATP and 0.1 GTP (pH 7.3 with KOH). All reagents were from Tocris, otherwise specified. mEPSCs were acquired at a 10 kHz sample frequency and filtered at 1/5 of the acquisition rate with a low-pass Bessel filter. The amplitude and frequency of the miniature excitatory events were calculated using a peak detector function with appropriate threshold amplitudes and areas using MiniAnalysis (Synaptosoft) and Prism (GraphPad Software Inc., La Jolla, CA). All experiments were performed at RT.

### Statistical analysis

Data are expressed as means ± SEM with individual experimental replications, as detailed in the figure legends. Normal distribution of data was assessed using the D’Agostino-Pearson’s normality test (n>6) or the Shapiro-Wilk test (n<6). The F-test was used to compare the variance between the two sample groups. The two-tailed unpaired Student’s t-test or the non-parametric Mann-Whitney’s U-test was used to compare two experimental groups based on data distribution. One-way or two-way ANOVA, followed by Tukey’s or Dunnett’s multiple comparison test, was used to compare more than two normally distributed experimental groups. The Kruskal-Wallis ANOVA was used to compare more than two non-normally distributed experimental groups, followed by Dunn’s multiple comparison test. The significance level was preset to p<0.05. Statistical analysis was done using Prism (GraphPad Software, Inc.).

## SUPPLEMENTARY MATERIALS

### SUPPLEMENTARY METHODS

#### Molecular Modeling of Wild-Type REST

The amino acid sequence V130-H267: “VAEDKCRSSKAKPFRCKPCQYEAESEEQFVHHIRIHSAK KFFVEESAEKQAKAWESGSSPAEEGEFSKGPIRCDRCGYNTNRYDHYMAHLKHHLRAGENERIYKCIICTYTTVSEYHWRKHLRNHFPRKVYTCSKCNYFSDRKNNYVQHVRTHTGERPYKCELCPYSSSQKTHLTRHMRTHSGEKPFKCDQCNYVASNQHEVTRHARQVHNGPKPLNCPHCDYKTADRSNFKKHVELHVNPRQFNCPVCDYAASKKCNLQYHFKSKH”, accounting for the first eight zinc finger domains (ZNF) of the wt mouse REST protein (mREST) was used as input for three independent structure-prediction program: D-I-TASSER^1,2^, AlphaFold2^3^, and RoseTTAFold^4^. Based on visual inspection with PyMOL^5^, the D-I-TASSER^1^ model was selected as the most reliable structural candidate for subsequent docking experiments (**Figure S7**).

#### Protein–DNA Docking

To assemble the REST–RE1 complex, the selected V130–H267 REST fragment and the RE1 DNA sequence GCTTTCAGCACCACGGACAGC (containing three upstream nucleotides) were submitted to HDOCK^6^, a hybrid docking server that combines template-based modeling with free docking. HDOCK^6^ uses a soft rigid-body docking strategy with limited backbone flexibility and relies on a knowledge-based scoring function integrating interface energy, buried surface area, and statistical potentials. The top-ranked binding mode (model 1, **Figure S6 - Step 4**) was used as the foundation for subsequent simulations.

#### Multiple Sequence Alignment

The REST amino acid sequence comprising residues V130–H267 was used as the query for a homology search against the Protein Data Bank (PDB)^7^. The NCBI Domain Enhanced Lookup Time Accelerated BLAST (DELTA-BLAST) algorithm^8^ was employed, with the maximum number of target sequences set to 50. An expectation value (E-value) threshold of 1e-5 and a word size of 3 were specified. Alignments were scored using the BLOSUM62 substitution matrix^9^, with gap penalties of 11 for gap existence and 1 for gap extension. Alignments were generated using a conditional compositional score matrix to correct for sequence-specific amino acid biases, and low-complexity regions were masked to minimize spurious matches driven by compositional heterogeneity.

#### Template Selection for REST-RE1 Complex

PDB entries retrieved from the multiple sequence alignment (MSA) were downloaded and inspected via PyMOL^5^. Duplicate chains and crystallization cofactors were removed, and records displaying missing residues were excluded both to prevent artifacts in the superimposition and to ensure structural completeness. Pairwise structural alignments were then performed between REST and each remaining protein chain, using REST as the mobile selection in the alignment procedure (**Figure S6 – Step 3; Figure S8A**). The corresponding RMSD values are reported in **Table S2**. Among the aligned structures, the protein–DNA complex deposited under PDB ID 1ubd was selected as the optimal template for REST–RE1 complex assembly (**Figure S8B**). This structure contains four zinc finger domains—only one fewer than REST—providing a biologically meaningful reference for modeling protein–DNA interactions. In addition, 1ubd contains no missing residues and yielded one of the lowest RMSD values in the dataset. Although a few entries produced marginally lower RMSDs, they failed to meet the full set of selection criteria outlined above. 1ubd was also instrumental in resolving ambiguities among the alternative docking poses by offering a validated spatial framework for the orientation of zinc finger domains relative to DNA. Furthermore, the conserved coordination geometry observed in 1ubd served as a structural guide for the accurate placement and chelation of Zn^2²^ ions within the modeled REST Zn-finger domains, ensuring correct folding and stability of the DNA-binding interface.

#### Metal3D Benchmarks

PDB entries 1a1h and 4r2a were selected as benchmark structures based on their low pairwise RMSD and their representation of distinct model organisms. Specifically, 1a1h corresponds to a murine zinc finger protein and 4r2a to a human zinc finger protein. Prior to analysis, all divalent zinc ions and nucleic acid molecules were stripped from the coordinate files, yielding protein-only structures suitable for being submitted to Metal3D convolutional neural network^10^. The coordinates of Zn^2+^ ions were then predicted at an isovalue probability threshold of *p = 0.9* and subsequently integrated into the corresponding protein-only structures. These reconstituted assemblies were superimposed onto the original PDB entries to quantify deviations between the experimental and predicted Zn^2+^ ion positions (**Figure S9**).

#### Zn^2²^ Ion Coordinate Prediction and Model Incorporation

The predicted REST protein structure was submitted to the Metal3D^10^ Colab Notebook, allowing the remaining convolutional neural network layers to initialize and process the input. A grid with a resolution of 0.5 Å was generated to compile per-residue metal-binding probability scores. Zn^2²^ 3D coordinates corresponding to an isovalue probability threshold of *p = 0.9* were then extracted and incorporated into the protein-only REST model. Multiple coordinate sets were predicted for each zinc-binding domain at this threshold. To select a single set of Zn^2²^ coordinates for each domain, the predictions were superimposed onto the zinc-binding site of the template structure (PDB: 1ubd) with PyMOL^5^ to assess positional deviations and identify the most structurally consistent coordinates (**Figure S10**).

#### Geometrical Optimization of ZNF Domains via Molecular Dynamics (MD) Simulations

The solution builder plug-in of the CHARMM-GUI web server^11,12^ was used to prepare the system and generate input files for MD simulations. The DNA–protein assembly was embedded in a rectangular box of TIP3P^13,14^ water molecules, with a minimum solute-to-box-edge distance of 15 Å applied along all directions. The total system charge was neutralized by adding Na^+^ and Cl^-^ ions until a physiological concentration of 0.15 M was reached. The whole system amounts to ∽112000 atoms. Simulations were performed with NAMD 2.14^15^ combining the AmberFf14SB^16^ force filed for the protein and the BSC1^17^ force field for the nucleic acid. Tetragonal periodic boundary conditions (PBC) were applied. The Particle Mesh Ewald method was used to treat electrostatic interactions and to avoid box surface effects with a maximum grid spacing of 1 Å and sixth-order beta splines. A cut-off of 9 Å was employed for Lennard-Jones interactions, which were smoothly switched off starting at 13 Å. Chemical bond distances involving hydrogen atoms were constrained using the SHAKE^18^ algorithm. A time-step of 2 fs was utilized. Prior to initiating the production phase, the system underwent a rigorous protocol consisting of energy minimization followed by a 250 ns-long equilibration, conducted in an isobaric–isothermal (NPT) ensemble (P = 1 atm, T = 310 K). Harmonic positional restraints were applied to the Cα atoms of the protein backbone to preserve the overall fold and the Colvar (collective variables) module^19^ was employed to maintain the proper zinc ion binding. Specifically, the Colvar Coordination Number variable was defined using a set of donor atoms surrounding the Zn^2²^ center to drive the system toward its native pseudo tetrahedral coordination geometry (**Figure S11**). Once the proper Zn^2²^ coordination environment was achieved, the simulation progressed to the production stage. At this point, all positional restraints on Cα atoms were released. The Coordination Number restraint on the zinc coordination shell was maintained for all production runs. Throughout the simulation, the target pressure (P=1) and temperature (T=310 K)were regulated using a Nosé–Hoover Langevin piston^20^ and Langevin thermostat^21^, ensuring stable thermodynamic conditions under NPT ensemble constraints.

## SUPPLEMENTARY TABLES

**Table S1.** Phospho-potential scores of putative mouse and human REST sites.

| Prediction software | REST S177/S182 |  | REST T294/T299 |  | REST S322/S327 |  | Notes |
| --- | --- | --- | --- | --- | --- | --- | --- |
|  | mouse | human | mouse | human | mouse | human |  |
| *Musite* | 0.260 | 0.260 | 0.414 | 0.414 | 0.782 | 0.779 | Generic PK |
| *NetPhos3.1* | 0.433 | 0.425 | 0.457 | 0.457 | 0.462 | 0.462 | CaMKs |
| *KinasePhos3* | NA | -- | NA | -- | NA | 0.790 | CMGC PKs |
| *Kinexus* | NA | No | NA | Yes | NA | Yes | Generic PK |
| *Scansite* | 1.183 | 1.253 | -- | -- | 2.067 | 2.067 | CLK2 |

The scores of the three putative CaMKIV phosphorylation sites obtained by scanning the mouse and human REST *consensus* sequences with distinct phosphor-site prediction softwares predicting the phosphorylation potential are shown. PK, protein kinase; CMGC, CMGC family kinases (Cyclin-dependent, Mitogen-activated, Glycogen synthase, and CDK-like kinases); CLK2, dual specificity protein kinase; --, not detected; ND, not available.

**Table S2.** RMSD values resulting from structural alignments.

| ***RCSBPDB ID*** | ***RMSD (Å)*** |
| --- | --- |
| *1a1f* | 1.49 |
| *1a1h* | 1.24 |
| *1a1i* | 1.19 |
| *1a1l* | 1.31 |
| *1jk1* | 1.18 |
| *1mey* | 1.47 |
| *1p47* | 1.23 |
| *1ubd* | 1.80 |
| *1zaa* | 1.29 |
| *2i13* | 1.93 |
| *2jp9* | 1.30 |
| *4r2a* | 1.23 |
| *5egb* | 2.75 |
| *5t0u* | 1.95 |
| *5v3g* | 2.69 |
| *5v3j* | 4.03 |
| *5v3m* | 4.05 |
| *5wjq* | 3.96 |
| *5yef* | 2.10 |
| *5yeg* | 8.41 |
| *6b0o* | 2.03 |
| *6blw* | 1.83 |
| *6e93* | 2.19 |
| *6ml2* | 1.36 |
| *6wmi* | 2.51 |
| *7eyi* | 3.04 |
| *7n5s* | 2.92 |
| *7n5w* | 10.52 |
| *7txc* | 6.72 |
| *7ysf* | 2.05 |
| *8e3e* | 3.25 |
| *8ssq* | 2.43 |
| *8sss* | 1.97 |
| *8sst* | 1.94 |
| *8ssu* | 2.61 |

RMSD values are in Å. Superimpositions were performed taking the protein chain of REST (V130-H267) as the mobile selection in PyMOL. RMSD values are given in Å.

## SUPPLEMENTARY FIGURES

**Figure S1.**
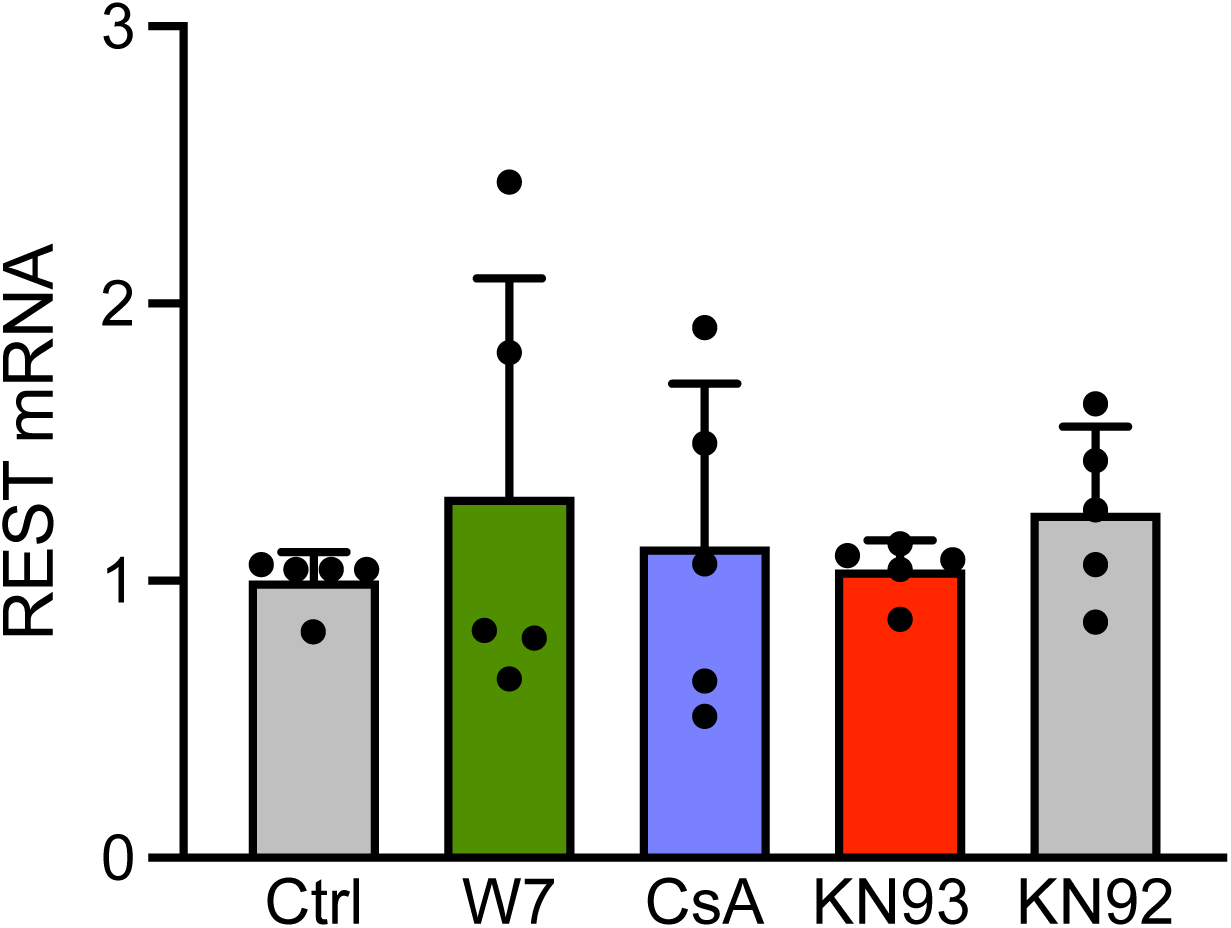
Inhibition of Ca^2+^-calmodulin-dependent activities does not affect REST transcription in primary cortical neurons. REST mRNA levels in primary cortical neurons (DIV14) treated with vehicle, the Ca^2+^-calmodulin antagonist W7 (20 µM), the phosphatase inhibitor cyclosporin A (CsA; 1 µM), the cell-permeable CaMK inhibitor KN93 (10 µM), and its inactive derivative KN92 (10 µM). Mean (± sem with individual experimental points) REST mRNA levels assessed by real-time qPCR. p>0.83, one-way ANOVA/Tukey’s tests (n=5 independent experiments).

**Figure S2.**
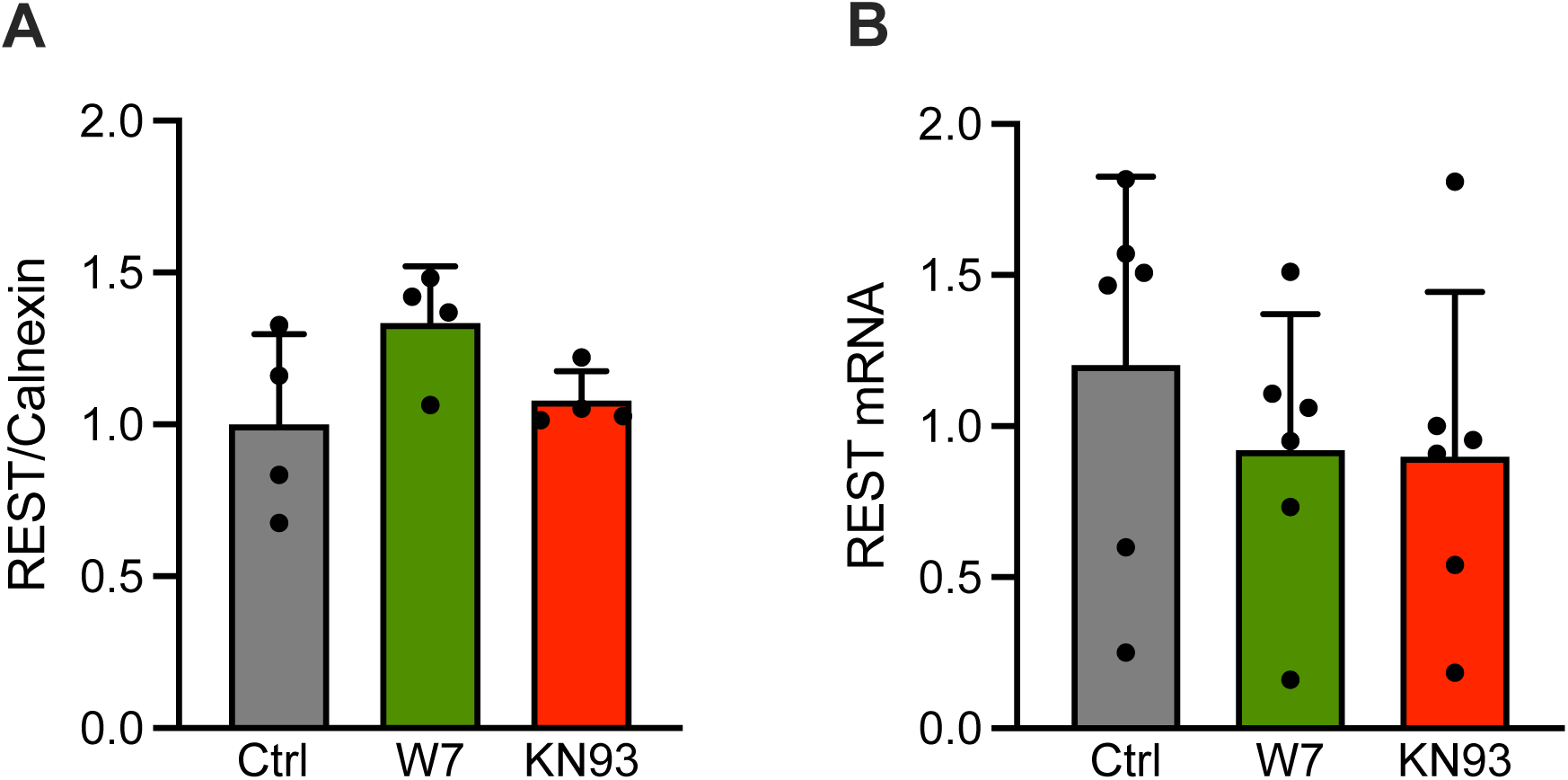
Inhibition of Ca^2+^-calmodulin-dependent activities does not affect REST transcription and translation in primary astrocytes. **A.** Immunoblotting of REST in primary astrocytes treated with vehicle, the Ca^2+^-calmodulin antagonist W7 (20 µM), and the CaMK inhibitor KN93 (10 µM). Calnexin was used as a control for equal loading. REST expression is shown as the mean ratio (± sem with individual experimental points) between REST and calnexin immunoreactivities. p>0.09; one-way ANOVA/Dunnett’s tests *vs* control (n=4 independent experiments). **B.** REST mRNA levels in primary astrocytes treated as in A. Mean (± sem with individual experimental points) REST mRNA levels assessed by real-time qPCR. p>0.54, one-way ANOVA/Dunnett’s tests *vs* control (n=6 independent experiments).

**Figure S3.**
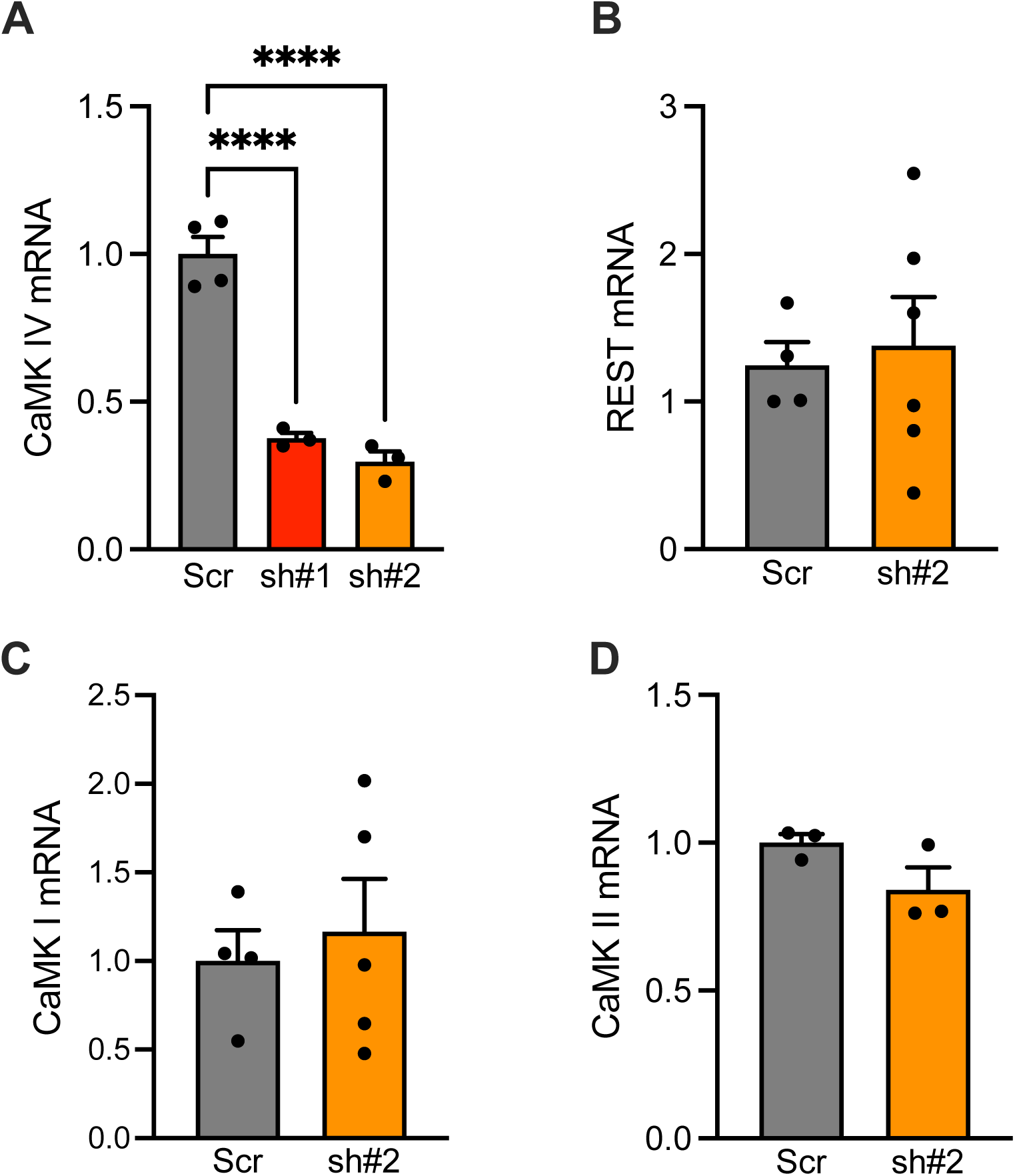
Silencing the CaMKIV by RNA interference does not affect REST, CaMKI, and CaMKII expression in primary neurons. Two short hairpin RNAs (sh#1 and sh#2) were used to test the efficacy of CaMKIV downregulation. The mean (± sem with individual experimental points) CaMKIV mRNA levels (**A**), assessed by real-time qPCR, were significantly downregulated by both shRNAs. CaMKIV downregulation by sh#2 did not affect the mRNA levels of REST (**B**), CaMKI (**C**), and CaMKII (**D**). A: ****p<0.0001, one-way ANOVA/Tukey’s tests (n=4 independent experiments). B-D: p=0.72 (B), p=0.65 (C), p=0.16 (D); unpaired Student’s *t*-test (n=6, 5, and 3 independent experiments for B, C, and D, respectively).

**Figure S4.**
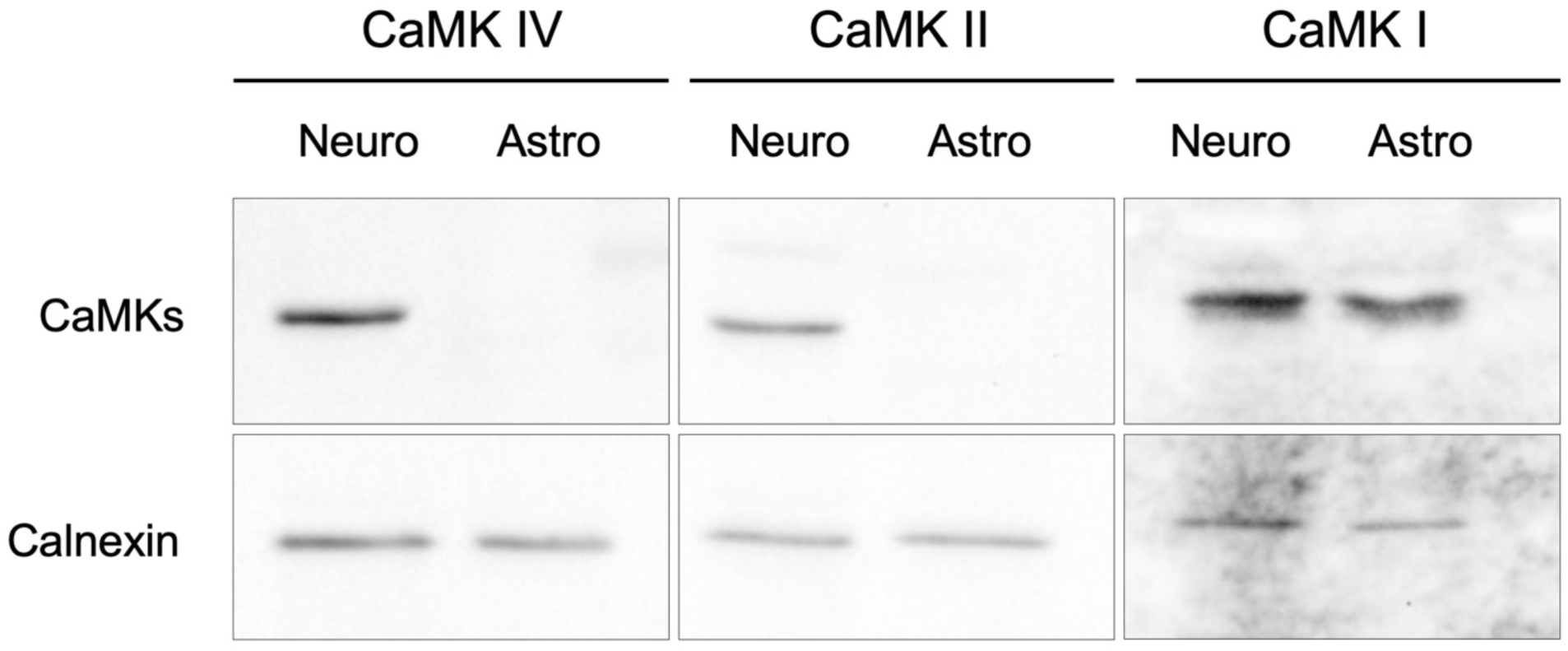
The expression of CaMKII and CaMKIV is neuron-specific. Immunoblots of CaMKI, CaMKII, and CaMKIV in primary neurons and primary astrocytes. Calnexin was used as a control for equal loading. While CaMKI is expressed to a similar extent in both neurons and astrocytes, CaMKII and CaMKIV display a purely neuron-specific expression.

**Figure S5.**
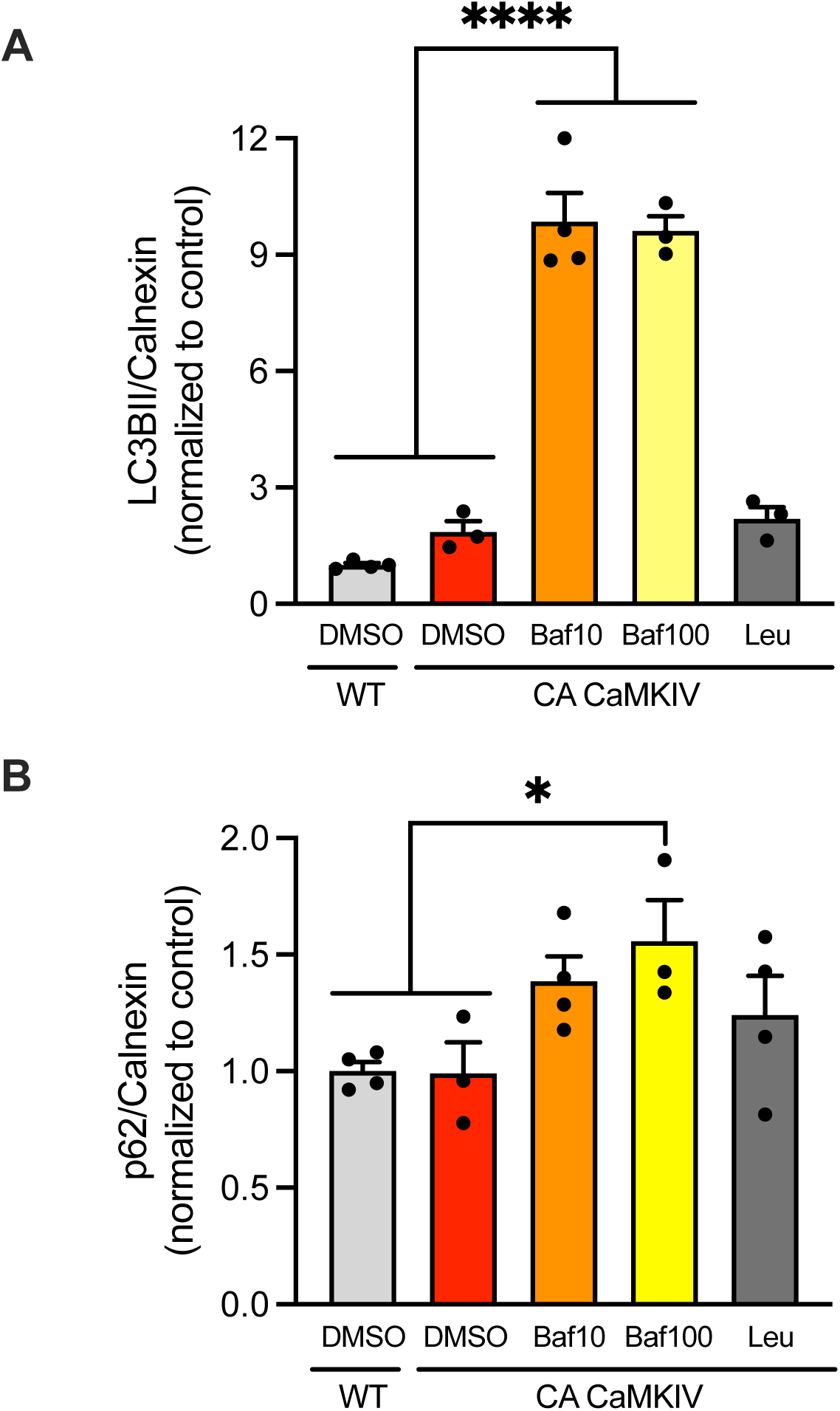
Bafilomycin induces autophagy blockade in Hek293T cells. Hek293T cells transfected with either WT or constitutively active CaMKIV (CA CaMKIV) and treated with cycloheximide, were treated for 16 h with vehicle (DMSO), bafilomycin (Baf, 10 and 100 nM), or leupeptin (Leu, 40 µM), as described in Figure 4C, which reports the representative immunoblots, monitoring the expression of REST, calnexin, and the autophagy markers p62 and LC3BII. Quantitative analysis of the autophagic block was performed by calculating the mean (± sem with individual experimental points) protein levels of the autophagy markers LC3BII (**A**) and p62 (**B**), shown as ratios with calnexin immunoreactivity under the various experimental conditions and normalized to the control (WT-REST/DMSO). A: ****p<0.0001 CA/Baf10 and CA/Baf100 *vs* control (n=3-6 independent preparations). B: *p=0.036 CA/Baf100 *vs* control (n=3-5 independent preparations). One-way ANOVA/Dunnett’s tests.

**Figure S6.**
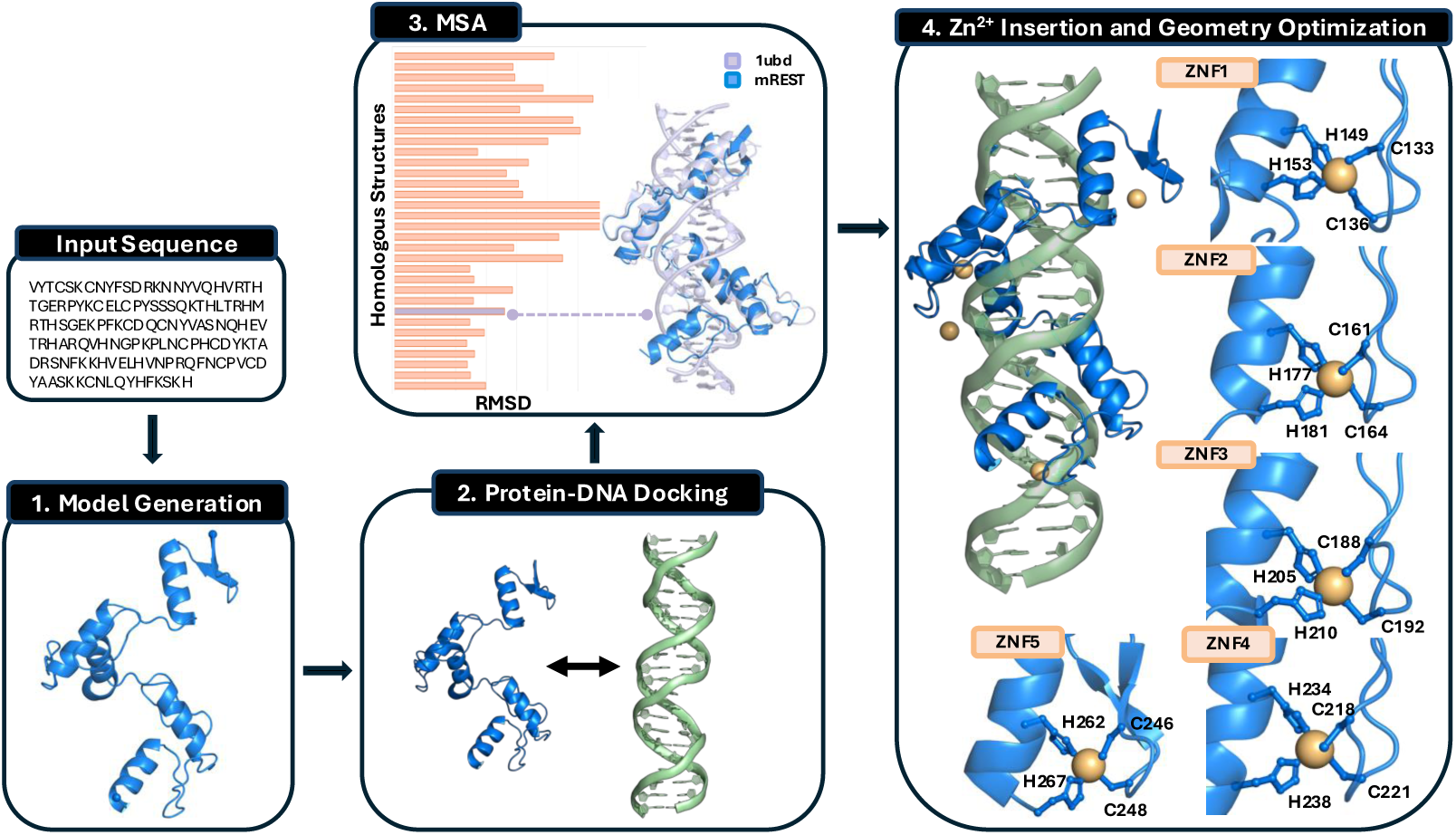
Schematic overview of the REST:RE1 structural modeling workflow. Schematic representation of the tailored, multi-step structural modeling pipeline developed to assemble the REST:RE1 protein–DNA complex. The workflow is organized into four interdependent modules, each enclosed in black-framed boxes and connected by arrows to indicate the sequential order of the steps. The pipeline begins with the generation of structural models for the individual REST and RE1 components (1), followed by a protein–DNA docking stage (2) to produce initial complex conformations. This is followed by multiple sequence alignment (3) aimed at identifying suitable structural templates to guide REST:RE1 interface refinement, helping to resolve ambiguities in docking poses, Zn^2+^ ions positioning, and to improve the biological relevance and accuracy of the modeled interactions. In the final step (4), ^2²^ ions are inserted into the Zn-finger domains of REST, and local coordination geometries are optimized to ensure correct chelation. Methodological details are provided in the Supplementary Information.

**Figure S7.**
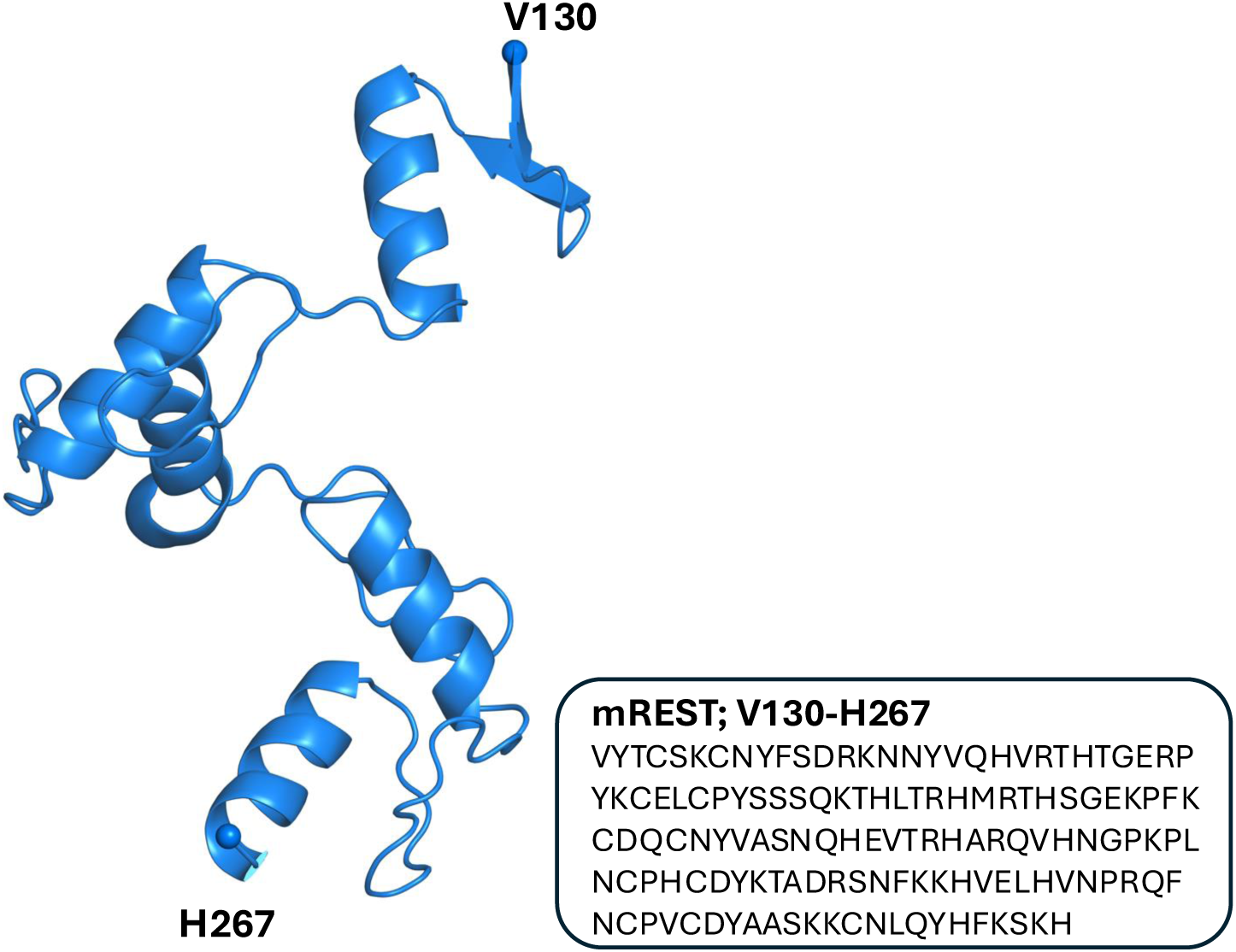
Molecular modeling of mREST ZNF domains. D-I-TASSER prediction for mREST amino acid string (V130-H267), displayed as blue cartoon; ***α***-carbons of N- and C-terminal residues are represented as spheres. In the black rectangle, the primary sequence employed as input to the D-I-TASSER structural modeling algorithm is shown.

**Figure S8.**
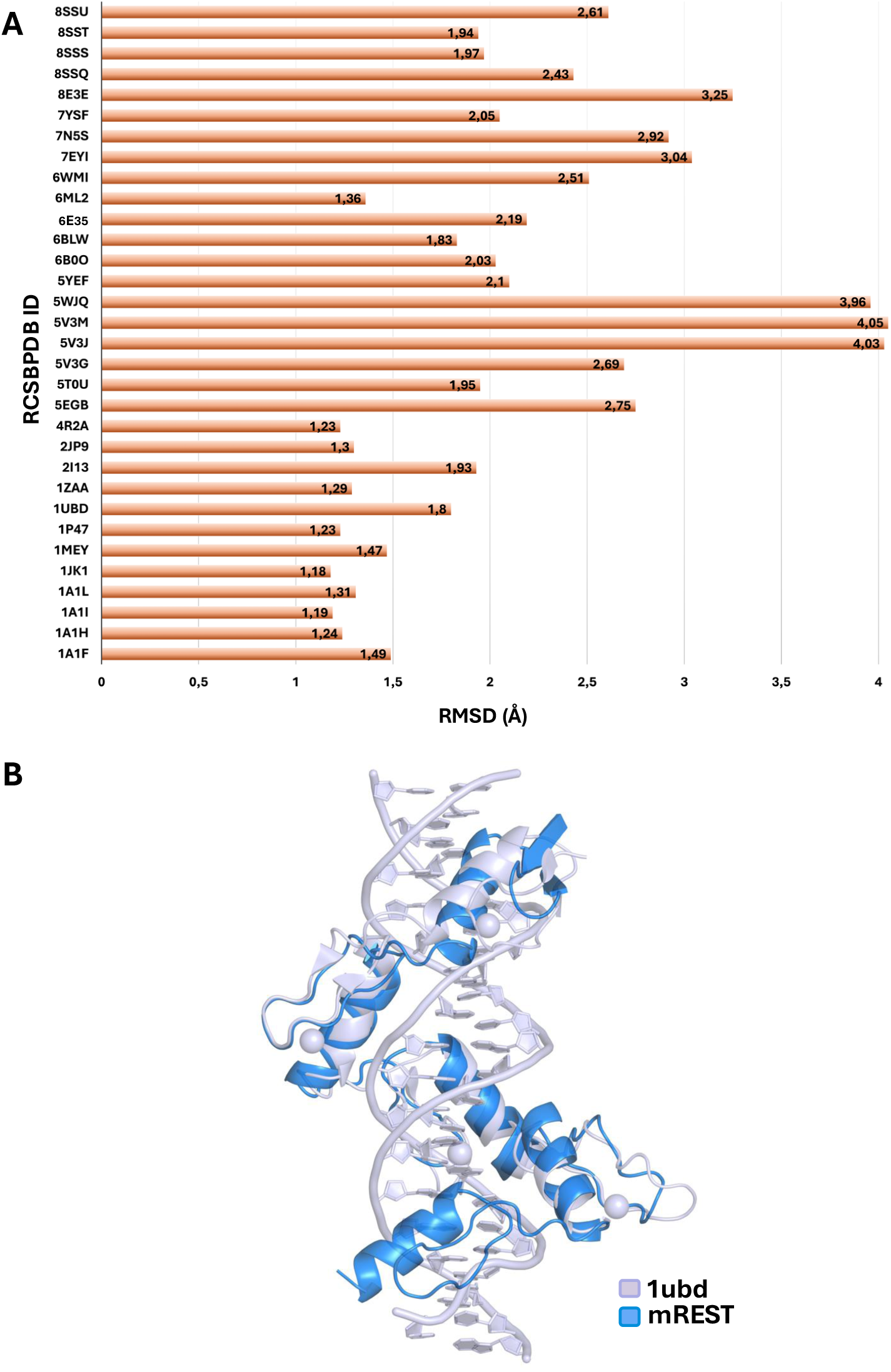
Structural template identification for REST:RE1. **A.** Root-mean-square deviations (RMSD, in Å) resulting from pairwise structural alignments between the modeled mREST ZF region and experimentally determined Zn-finger domains from the RCSB Protein Data Bank. Individual RMSD values for each alignment are annotated at the end of each bar. **B.** Superimposed structures of mREST (blue) and PDBID 1ubd, in lilac. Both protein and DNA components are displayed in cartoon representation. Coordinated Zn^2²^ ions are shown as spheres.

**Figure S9.**
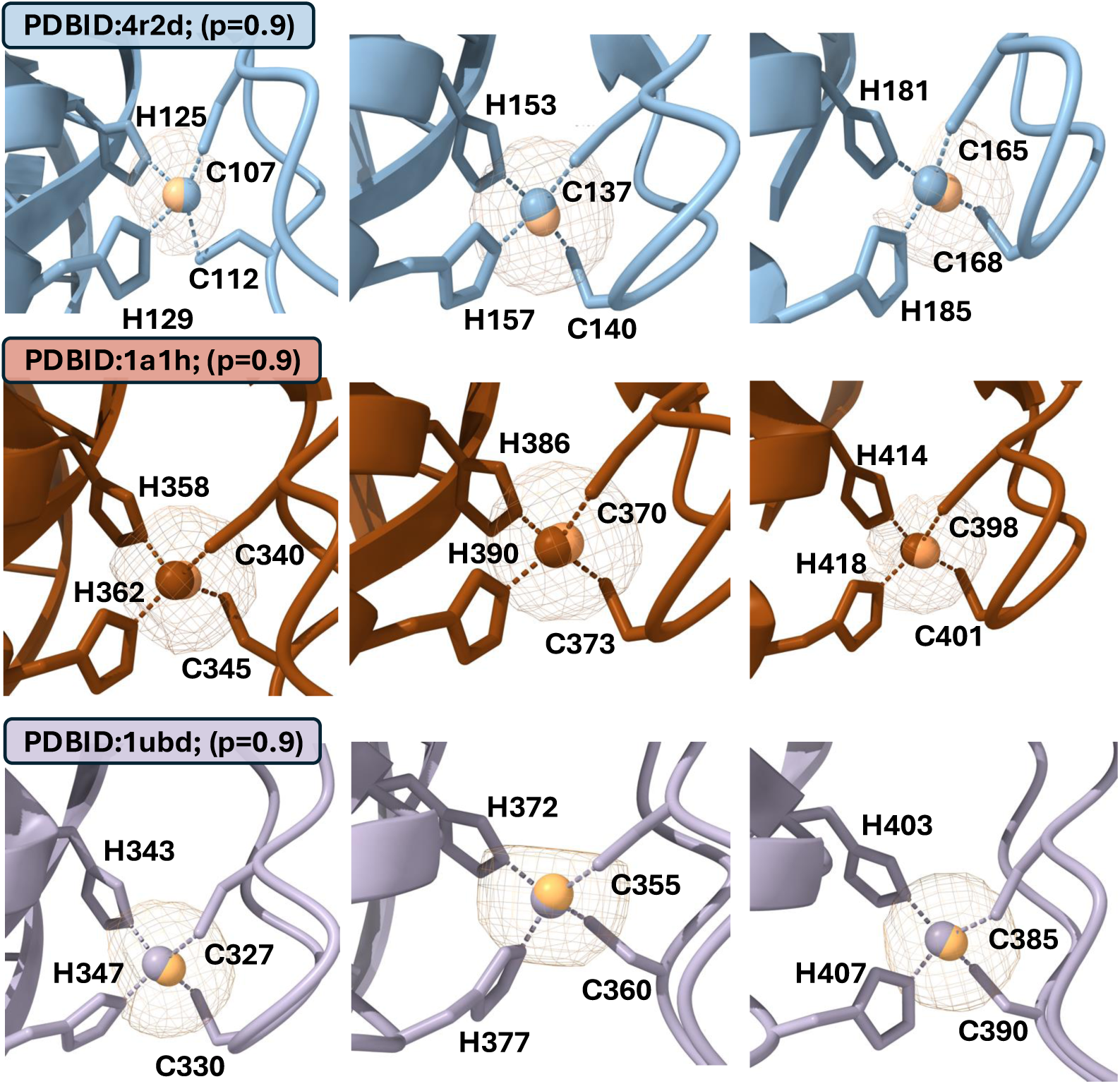
Benchmarking of Metal3D predictions for Zn^2+^ coordination sites. Comparison between predicted (orange) and known Zn^2+^ position for a set of experimentally determined protein structures. Protein backbones are reported in cartoon, while the side chains of zinc-chelating residues are shown as sticks. Transparent mesh surfaces indicate probability iso-surfaces of *p* = 0.9 for ion position.

**Figure S10.**
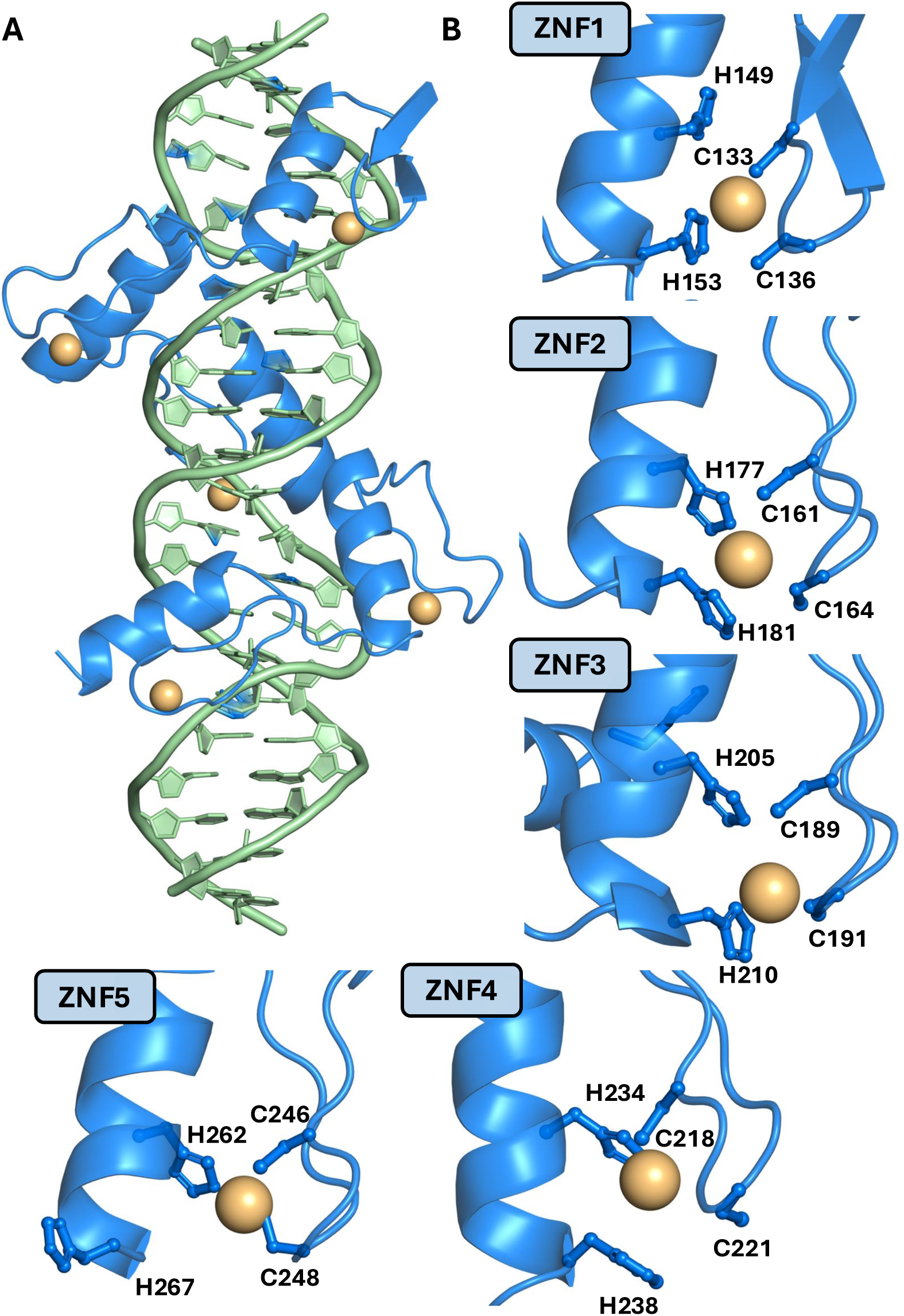
Pre-optimized REST ZNF domains docked onto the RE1 DNA element. **A.** Top-ranked docking conformation of the mREST ZNF array bound to its cognate RE1 DNA motif. **B.** Pre-optimized structures showing non-refined geometry of ZNF1 through ZNF5. Protein and DNA molecules are depicted in cartoon representation and colored blue and green, respectively; Zn^2+^-coordinating side chains are rendered as ball-and-stick, while coordinated Zn^2²^ ions are shown as orange spheres.

**Figure S11.**
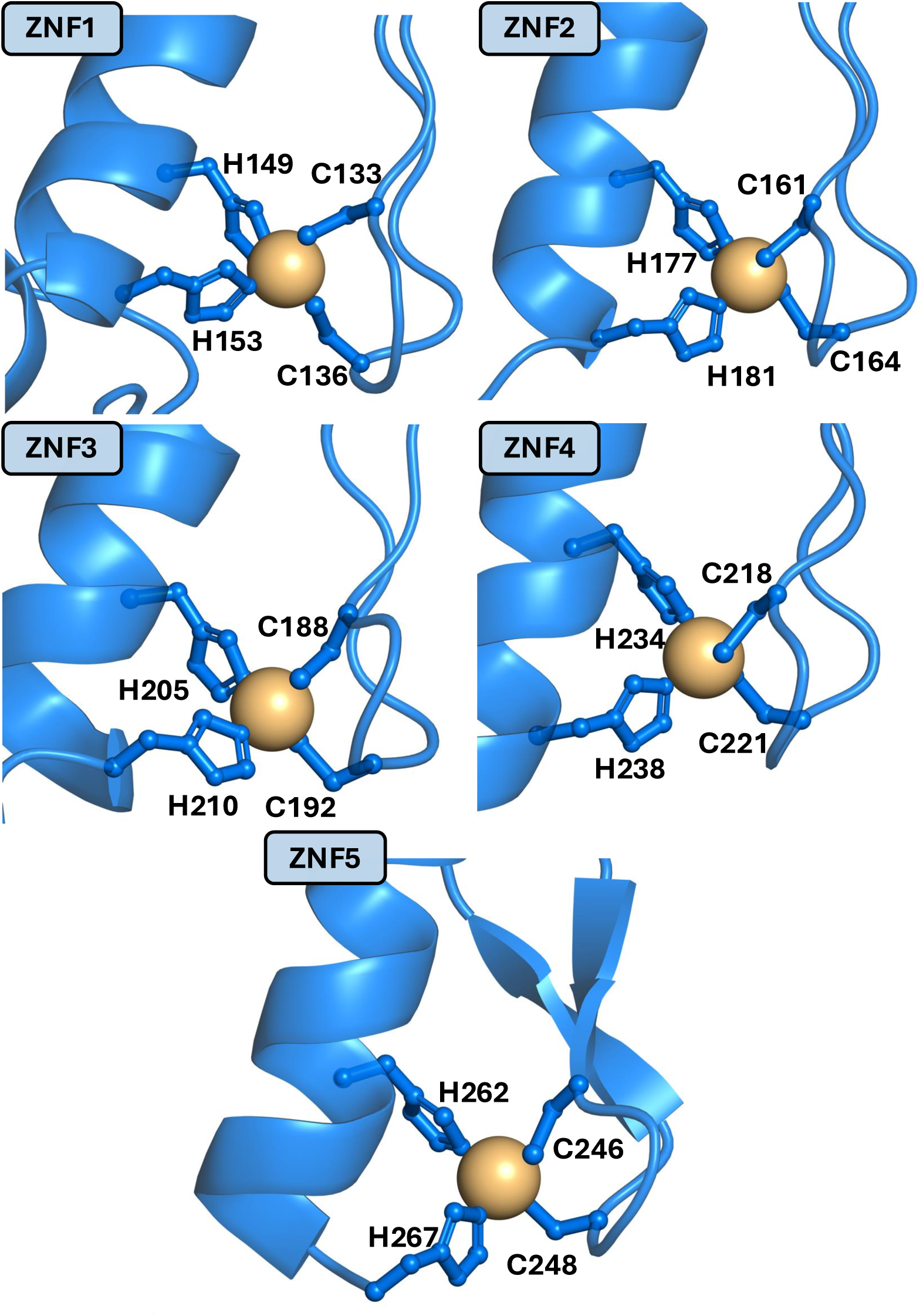
Optimized Zn^2+^-coordination spheres. Representative snapshots at the end of the refinement step, showing canonical pseudo-tetrahedral coordination geometries of all ^2²^ ions. The protein backbone is shown as blue cartoon, with coordinating residues represented as sticks and spheres. Zn^2²^ ions are visualized as orange spheres.

**Figure S12.**
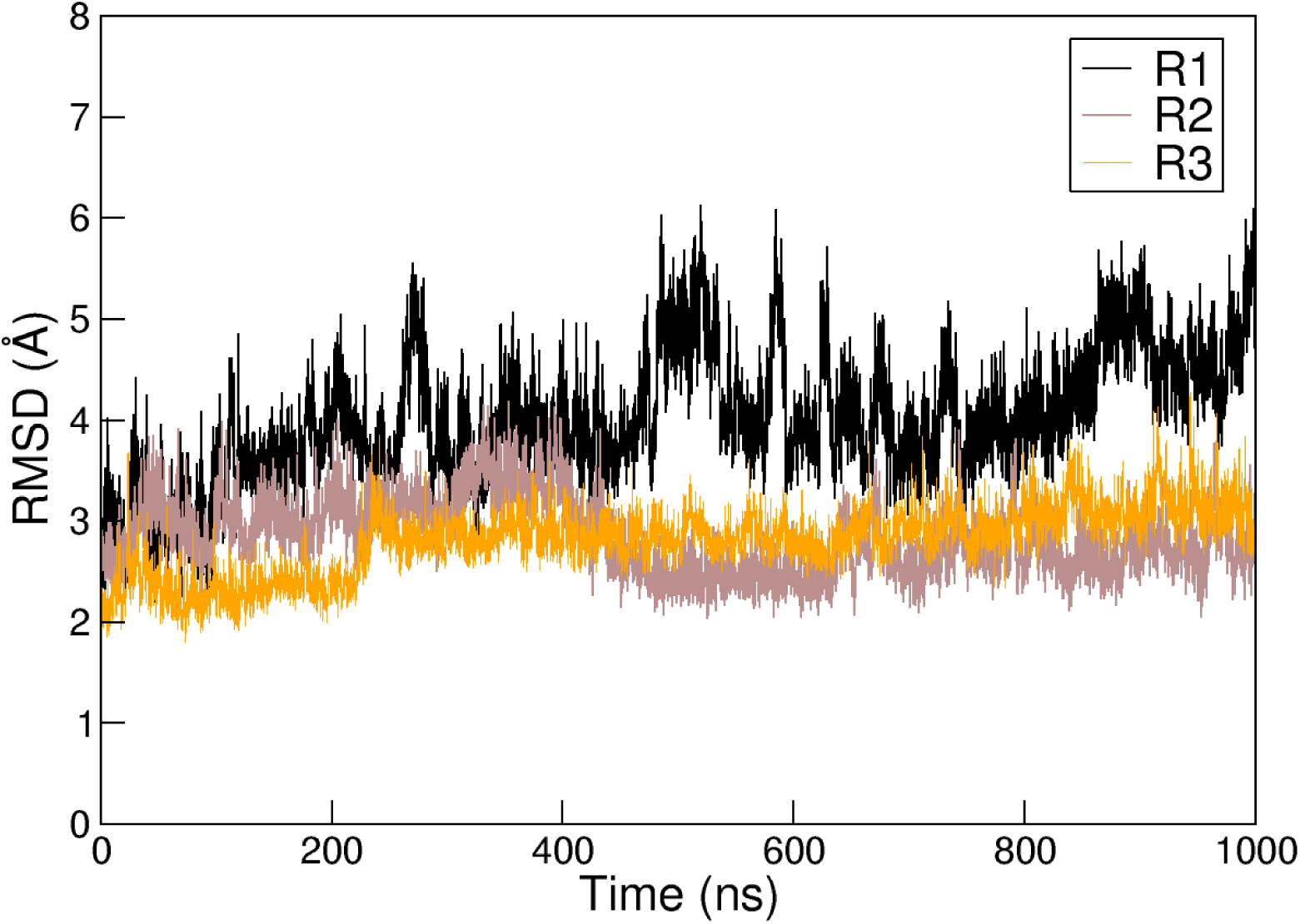
RMSD profiles of the WT mREST:RE1 complex. Time evolution of the root mean square deviation (RMSD) profiles of the full mREST:RE1 complex calculated over the course of three independent MD simulations. Each trajectory was generated under identical simulation conditions, starting from the same minimized and equilibrated initial structure. RMSD values were computed using all heavy atoms of both mREST and the RE1 DNA segment, after least-squares fitting to the initial frame based on the complete assembly.

**Figure S13.**
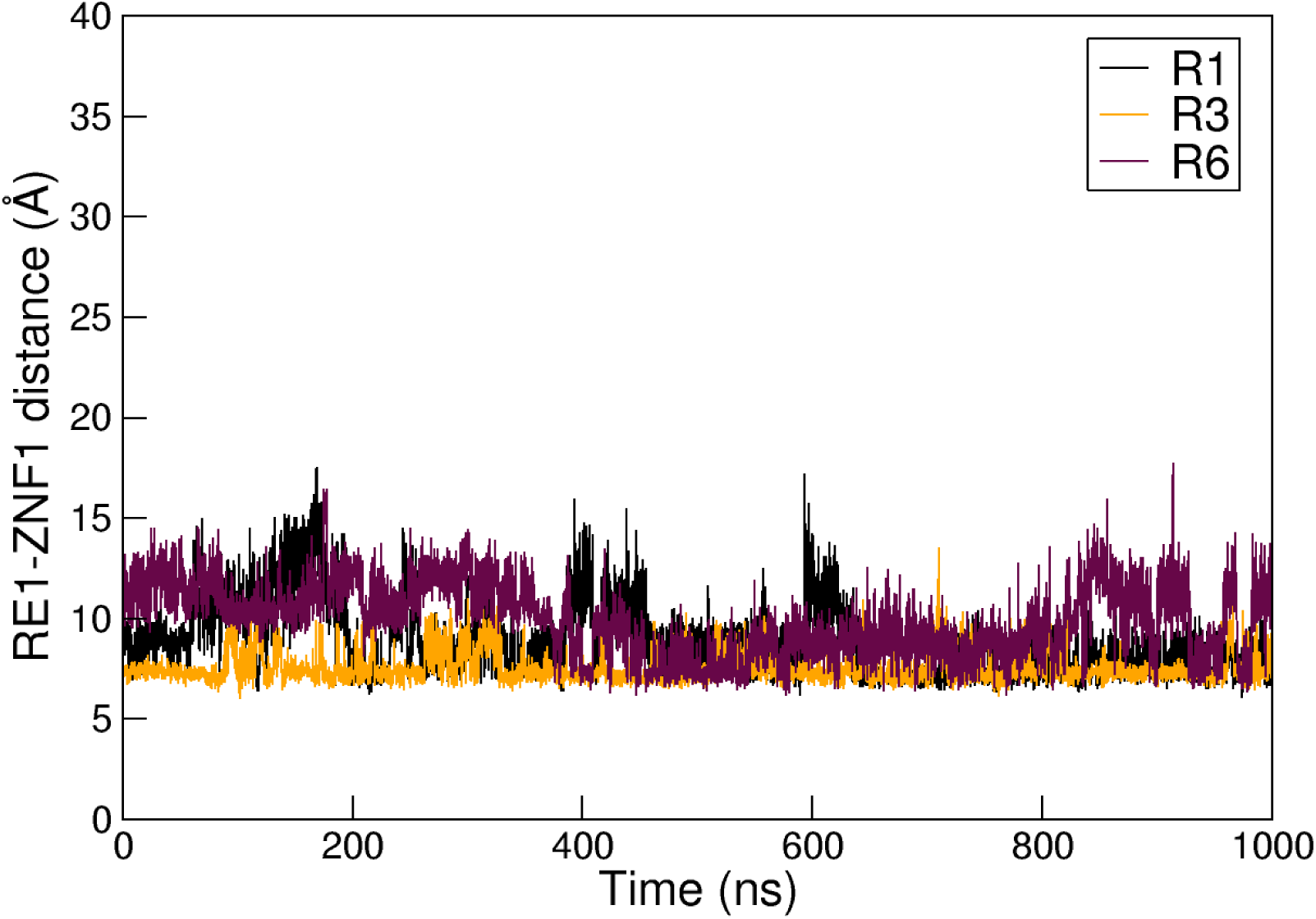
Time evolution of the RE1-ZNF1 distance for selected replicas of mutant S322E. Distance between RE1 and ZNF1 measured over time for three independent MD simulations, replica R1 (black), R3 (orange), and R6 (maroon), of the S322E phosphomimic variant of the mREST:RE1 complex. The distance is calculated between the phosphorus atom of the RE1 DNA segment and the center of mass of the Zn^2+^-coordinating atoms within the first ZNF domain (ZNF1) of mREST.

**Figure S14.**
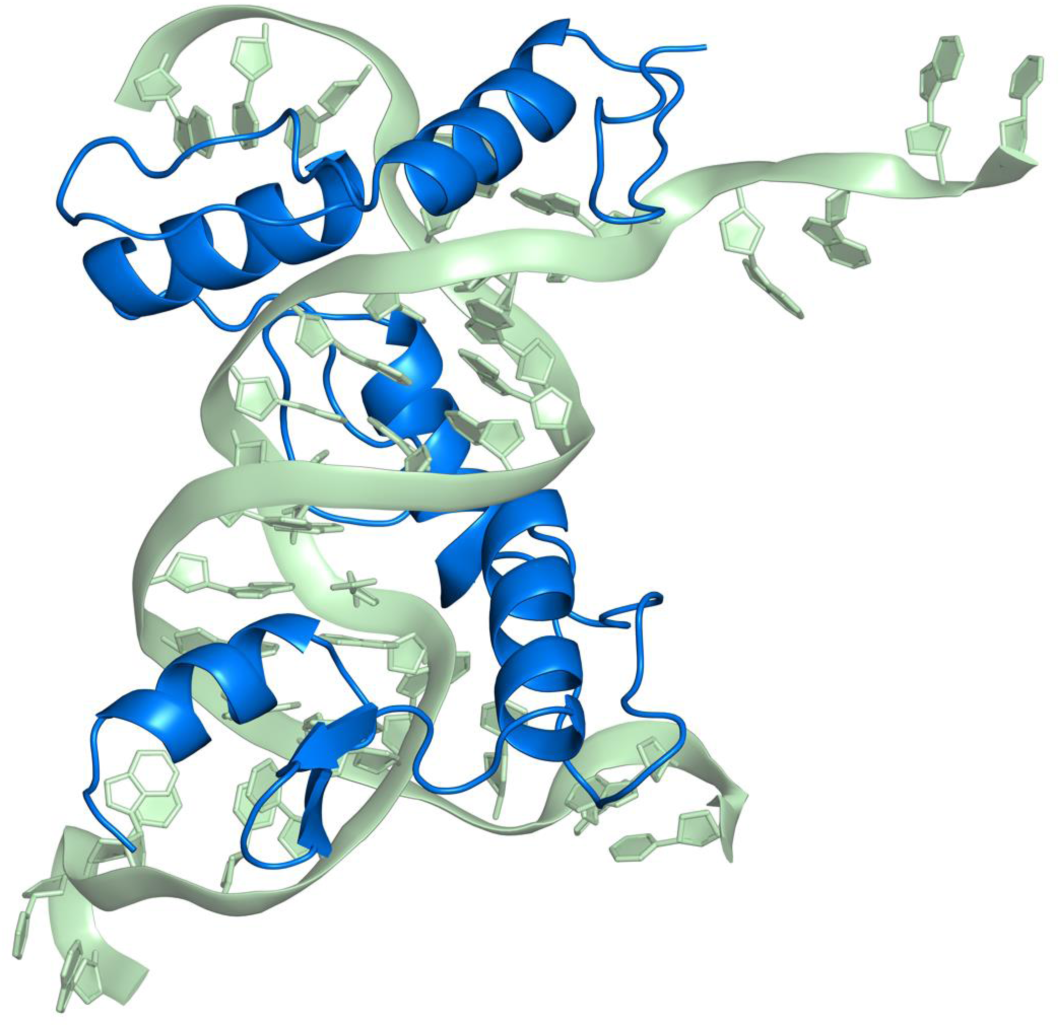
Fragilization of DNA termini leads to S322E system collapse. Structural snapshot illustrating the collapse of the REST:RE1 complex carrying the S322E phosphomimic substitution. The protein is shown in blue and the DNA double helix in green. The image captures a severe distortion of the canonical B-form DNA conformation, with pronounced bending and fraying at the terminal base pairs, leading to global destabilization of the protein–DNA interface. The REST domain loses its defined binding orientation, with helices disengaging from the DNA major groove and extensive disruption of side-chain contacts. These conformational aberrations underscore the critical importance of DNA end integrity in maintaining REST:RE1 structural stability and suggest that perturbations such as the S322E mutation may exacerbate interface vulnerability by promoting electrostatic repulsion and misalignment.

**Figure S15.**
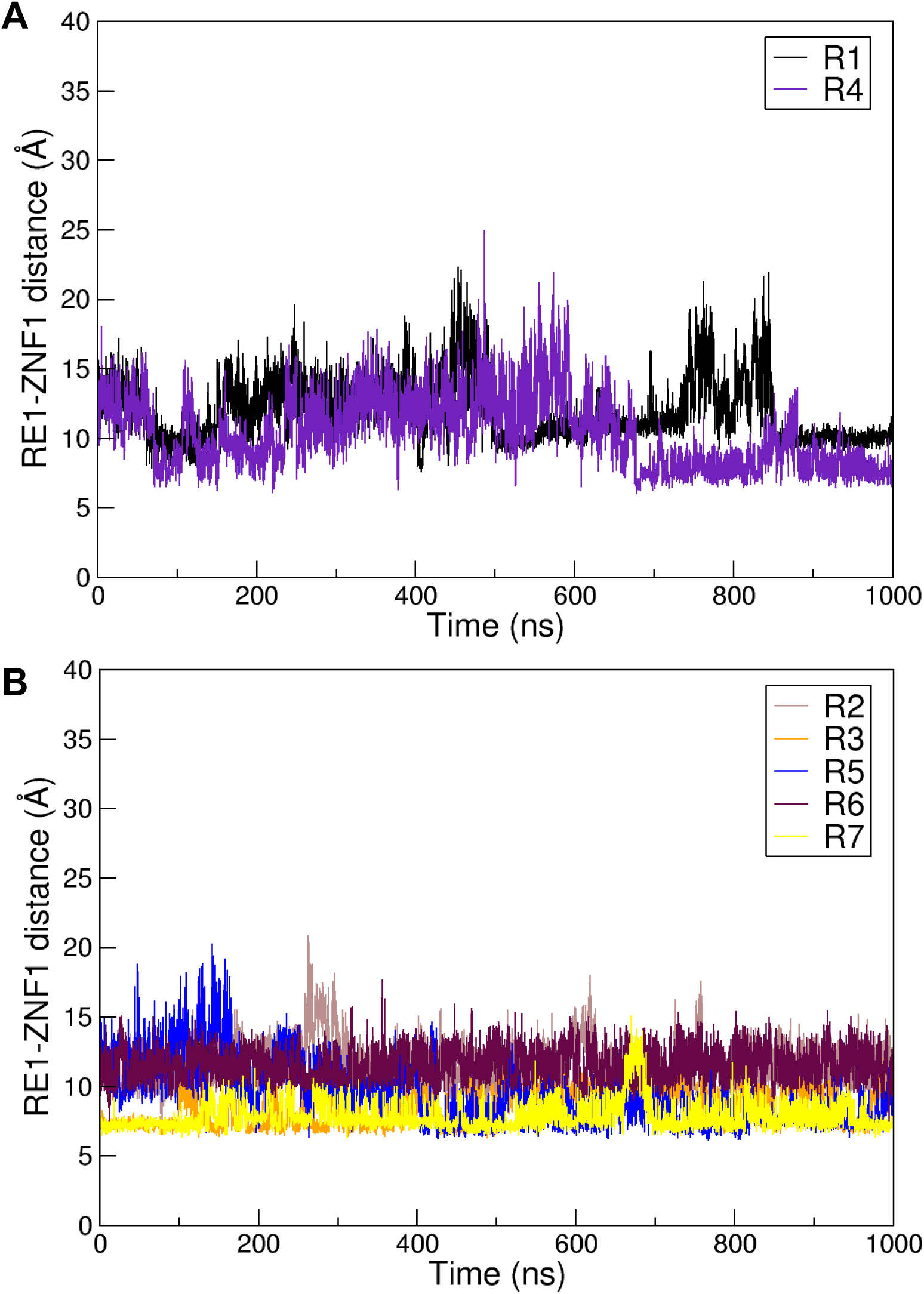
Time evolution of the RE1–ZNF1 distance for selected replicas of mutant T294E. **A.** Distance between RE1 and ZNF1measured over time for two independent MD simulations, replica R1 (black) and R4 (purple) of the T294E phosphomimic variant of the mREST:RE1 complex. **B.** Same for replica R2 (orange), R3 (brown), R5 (blue), R6 (maroon), and R7 (yellow).

## Notes

### Competing Interest Statement

The authors have declared no competing interest.

### Summary of Updates

Acknowledgement of additional funding and shorter summary.

